# MOSAIC: A Structured Multi-level Framework for Probabilistic and Interpretable Cell-type Annotation

**DOI:** 10.64898/2026.01.31.702798

**Authors:** Mingyu Yang, Jing Qi, Minwen Lan, Jiacheng Huang, Shuilin Jin

## Abstract

Accurate cell-type annotation is a foundational task in single-cell RNA sequencing analysis, yet remains fundamentally challenged by cellular heterogeneity, gradual lineage transitions, and technical noise. As single-cell atlases expand in scale and resolution, most existing annotation approaches operate at a single analytical level and encode cell identity as fixed categorical labels, limiting their ability to represent uncertainty, mixed biological states, and population-level structure. Here we introduce MOSAIC (Multi-level prObabilistic and Structured Adaptive IdentifiCation), a structured multi-level annotation framework that integrates cell-level marker evidence with cluster-level population context within a unified probabilistic system. Rather than treating annotation as an independent per-cell prediction task, MOSAIC formulates cell-type assignment as a coordinated multi-level inference process, in which probabilistic evidence at the single-cell level is aggregated, constrained, and refined by population context. MOSAIC integrates direction-aware marker scoring with dual-layer probabilistic representation and adaptive cross-level refinement, enabling uncertainty to be quantified and propagated across biological scales. This design yields coherent annotations that preserve fine-grained single-cell variation while maintaining population-level consistency, and allows ambiguous or transitional states to be represented explicitly rather than collapsed into hard labels. Across six diverse tissues and under controlled dropout perturbations, MOSAIC consistently matches or outperforms representative marker-based, reference-based, and machine-learning annotation methods. Beyond accuracy, MOSAIC provides structured uncertainty estimates and coherent population-level structure, enabling the identification of stable intermediate cell states that arise from gradual lineage transitions rather than technical noise. Together, MOSAIC advances cell-type annotation from a single-level classification task to a structured multi-level inference problem, and establishes a general, interpretable, and uncertainty-aware computational framework for large-scale single-cell analysis.

## Introduction

Single-cell RNA sequencing (scRNA-seq) has enabled high-resolution characterization of cellular heterogeneity across tissues, developmental trajectories, and disease states^1,2^. As single-cell atlases continue to expand in scale and granularity, cell-type annotation has emerged not merely as a labeling task, but as a core computational problem involving uncertainty, hierarchy, and evidence integration^3,4^. Contemporary analyses increasingly require principled representations of ambiguous and transitional states, quantification of confidence rather than hard assignments, and coherence between single-cell observations and population-level organization^5,6,7,8,9^.

A wide range of annotation strategies has been developed, yet most capture only partial aspects of this complexity^2,10^. Marker-based approaches, including systems such as scType^11^, which formalized the joint use of positive and negative markers, offer transparent, gene-centric reasoning at the single-cell level. However, they typically rely on deterministic scores or hard assignments and lack mechanisms to propagate uncertainty, integrate population-level structure, or support adaptive refinement across biological scales^2,5,10,12^. Reference-based methods, such as SingleR^13^, can perform well when query data closely match curated atlases, but their accuracy degrades under domain mismatch or incomplete reference coverage^14^.

Large-scale machine-learning and deep-learning classifiers (e.g., CellTypist^15^, scDeepSort^16^) enable high-throughput prediction but ultimately encode cell identity as discrete categorical outcomes. Cluster-level aggregation or rule-based systems partially stabilize predictions by leveraging population structure, but remain unable to systematically represent intra-cluster heterogeneity or mixed identities^10,12,17^.

These limitations reflect a shared methodological constraint: most existing approaches operate at a single analytical resolution^2,18,19^. They do not explicitly model how marker evidence at the single-cell level accumulates into coherent population-level structure, nor how population-level signals should, in turn, guide refined decisions for individual cells^12,20,21,22^.

From a methodological perspective, these limitations point to a deeper issue: cell-type annotation is most often formulated as an independent per-cell classification task, rather than as a structured inference problem that explicitly couples single-cell evidence with population-level organization. In such formulations, gradual differentiation continua, ambiguous populations, and mixed phenotypes are forced into fixed labels, obscuring uncertainty and conflating biologically meaningful intermediate states with low-confidence assignments^5,6,7,8,9^. Importantly, such ambiguity often reflects genuine intermediate or transitional biological states, particularly in immune systems, rather than simply reflecting annotation errors^6,7,8,23^.

To address this conceptual gap, we introduce MOSAIC (Multi-level prObabilistic and Structured Adaptive IdentifiCation), a structured multi-level probabilistic framework for cell-type annotation that jointly integrates marker evidence, probabilistic inference, and population structure across biological scales (Fig. 1). Rather than treating cells and clusters as independent annotation units, MOSAIC performs coordinated reasoning through two complementary information paths. At the cell level, direction-aware marker scoring uses positive and negative markers to generate probabilistic identity vectors that capture lineage evidence and uncertainty. At the population level, these vectors are aggregated within clusters to construct soft consensus profiles and relative enrichment, enabling systematic discrimination between confidently supported identities, mixed lineages, and weakly supported states.

**Figure 1.**
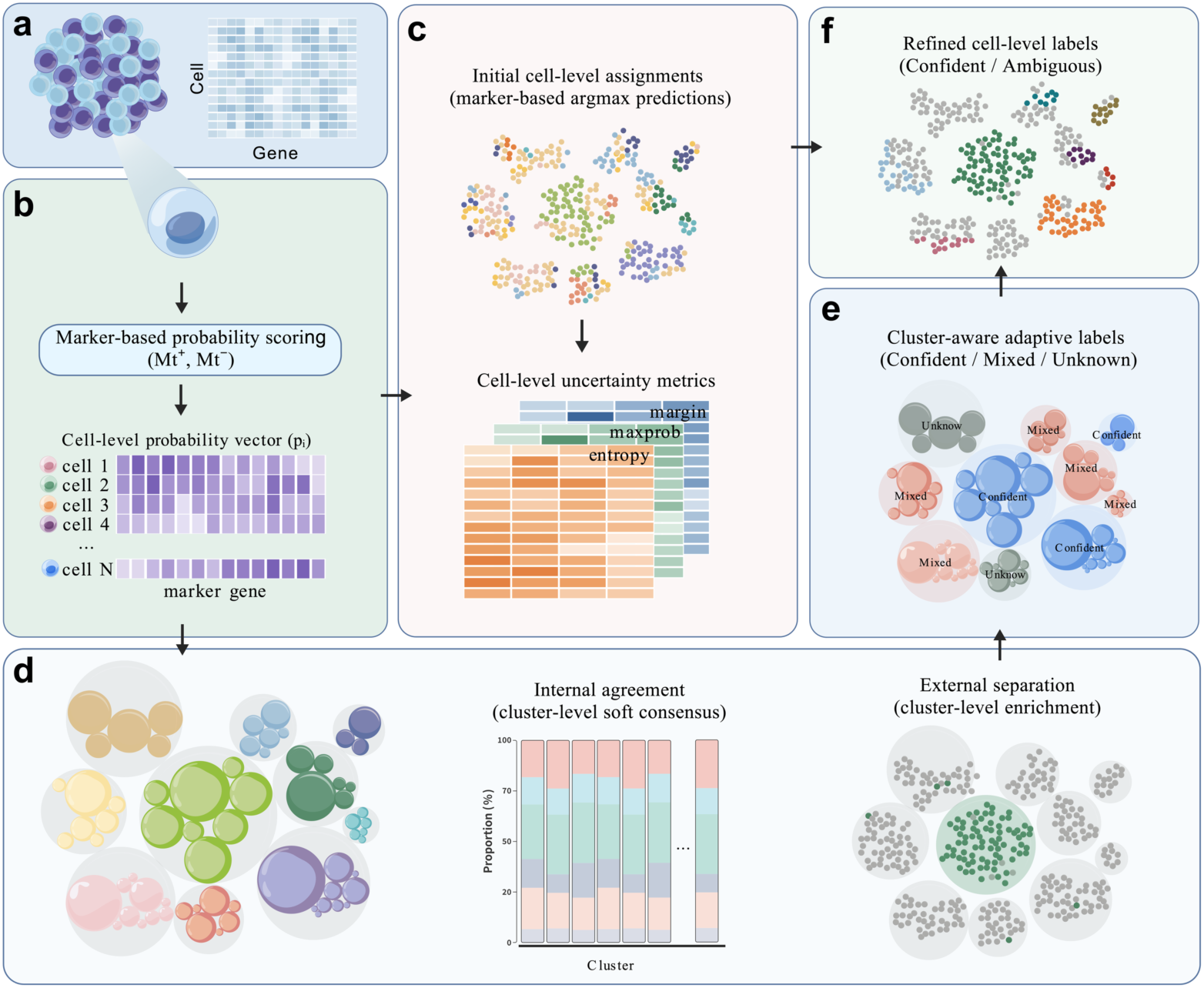
Overview of MOSAIC: a structured multi-level inference framework for probabilistic and interpretable cell-type annotation. **(a)** Single-cell expression profiles provide the input data, represented as a cell-by-gene matrix capturing heterogeneous marker expression across individual cells. **(b)** Direction-aware marker scoring integrates positive and negative markers to generate a full posterior probability vector for each cell, encoding lineage evidence and uncertainty rather than producing hard assignments. **(c)** Cell-level probabilistic representations are summarized for diagnostic visualization using maximum a posteriori (argmax) labels, together with complementary uncertainty measures, including maximum class probability, probability margin, and normalized entropy. **(d)** Cell-level probability distributions are aggregated at the cluster level to capture population-level structure through complementary views. Left, clusters are represented by outer grey circles with area proportional to cluster size, while inner colored circles indicate candidate identities supported within each cluster; circle size reflects relative probability mass, with the dominant soft consensus identity (soft1) and secondary identities explicitly represented. Middle, internal agreement quantifies intra-cluster coherence by aggregating cell-level probability distributions across identities. Right, external separation measures the relative enrichment of candidate identities across clusters, distinguishing cluster-specific signals from broadly distributed identities. **(e)** A cluster-aware adaptive decision module integrates cluster-level internal agreement and external separation, together with overall support strength, to classify clusters as Confident, Mixed (A/B), or Unknown, explicitly encoding ambiguity at the population level. **(f)** Final refined cell-level annotations are obtained by reconciling cluster-level decisions with local cell-level probabilistic evidence, assigning labels only where both levels are consistent and retaining ambiguous cells otherwise. Together, MOSAIC formulates cell-type annotation as a coordinated multi-level inference process, in which uncertainty is a first-class representation and population structure actively constrains single-cell interpretation, rather than being applied as a post hoc correction.

These two layers are unified through an adaptive refinement module that enforces consistency between local cell-level evidence and global population-level structure. The resulting annotations preserve fine-grained single-cell variation while maintaining cluster-level coherence, yielding a mosaic view of cellular identity in which ambiguous and transitional states are explicitly modeled rather than suppressed. Importantly, this integration does not rely on iterative feedback or heuristic post hoc corrections; instead, it formulates annotation as a structured probabilistic inference problem that synthesizes complementary evidence streams within a unified framework.

We evaluate MOSAIC across six diverse tissues—PBMC, liver, colon, pancreas, lung, and retina, as well as under controlled dropout perturbations that progressively degrade marker signal. Across these settings, MOSAIC demonstrates robust performance while simultaneously providing interpretability, uncertainty-aware annotations that remain coherent across scales. Together, these results position MOSAIC as a general computational framework for transparent, structured and probabilistic cell-type annotation in large-scale single-cell studies.

## Results

### Result 1 | A structured multi-level annotation framework integrating probabilistic cell-level evidence and population-level consensus

To illustrate the core inference structure of MOSAIC, we use the PBMC dataset as a compact and well-characterized example in which canonical immune populations, closely related states, and transitional phenotypes coexist. PBMC serves here as an explanatory system for visualizing how probabilistic evidence is generated, aggregated, and reconciled across analytical levels within a single annotation framework. The same four-stage procedure was applied to all other tissues analyzed in this study; PBMC is shown here for clarity and interpretability.

For reference, curated cell-type labels were overlaid on the UMAP embedding to visualize the global organization of major immune compartments, including T cells, B cells, monocytes, dendritic cells, and natural killer cells (Fig. 2a, upper left). These labels were used exclusively for visualization and evaluation and were not incorporated into any stage of the MOSAIC inference process.

**Figure 2.**
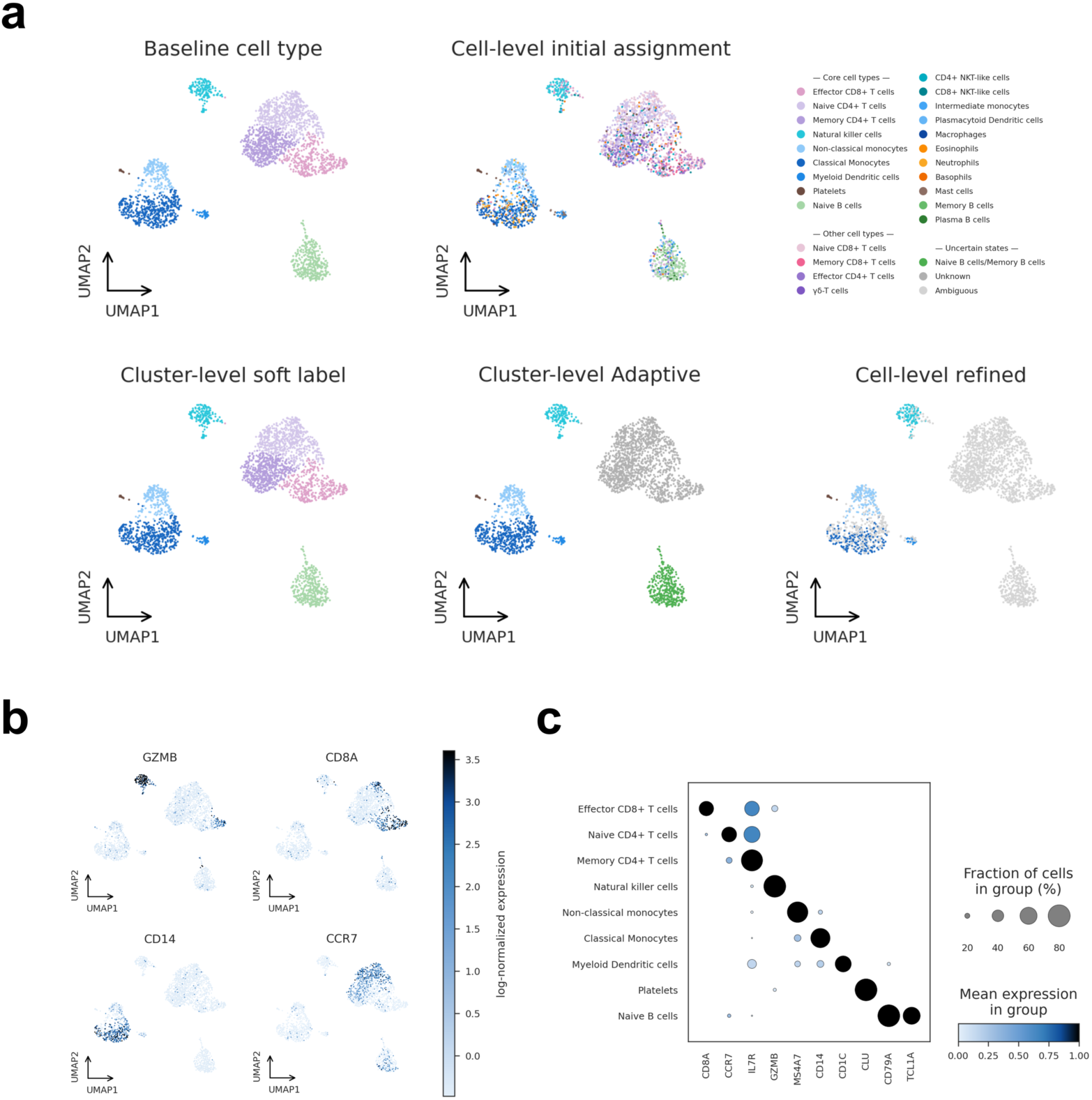
Multi-level probabilistic annotation behavior of MOSAIC illustrated in PBMC. **(a)** UMAP visualization of PBMC under successive stages of multi-level annotation. Upper left, curated baseline cell-type labels shown solely as a biological reference for visualization and evaluation; these labels are not used in any stage of MOSAIC inference. Upper right, initial cell-level assignments obtained by maximum a posteriori (argmax) decoding of marker-based probability vectors, which recover major immune populations but exhibit substantial mixing between closely related states. Lower left, cluster-level soft identities derived from aggregation of full cell-level probability distributions within Leiden clusters, resulting in sharpened population structure and suppression of scattered misassignments. Lower middle, adaptive cluster-level decisions classifying clusters as Confident, Mixed, or Unknown based on probability strength, separation, and relative enrichment, with ambiguous clusters collapsed into a neutral background to explicitly encode population-level uncertainty. Lower right, refined cell-level annotations obtained by reconciling cluster-level decisions with local probabilistic evidence, assigning labels only where both levels are consistent and retaining ambiguity otherwise. **(b)** UMAP overlays of representative canonical lineage markers (CCR7, CD8A, CD14, and GZMB) demonstrate spatially coherent and lineage-specific expression patterns that align with adaptively inferred cluster identities. **(c)** Dot-plot summary of marker expression aggregated by adaptive cluster labels, showing both mean expression and the fraction of expressing cells across immune compartments, confirming that inferred identities are supported by canonical marker signatures. PBMC is shown as a representative explanatory example; the same multi-level procedure was applied uniformly across all other tissues analyzed in this study. For visualization of refined cell-level annotations in PBMC only, an auxiliary probability threshold (*θ*_*vis*_ = 0.1) was applied to UMAP embeddings to improve visual interpretability of low-confidence and transitional cells. This threshold was used exclusively for visualization and did not affect any annotation, refinement, or quantitative evaluation, all of which relied on the default parameters described in the Methods.

MOSAIC begins by transforming the expression matrix into probabilistic cell-level evidence. Direction-aware marker scoring jointly integrates positive and negative marker information to assign each cell a full probability vector over candidate identities. Taking the maximum a posteriori assignment recovers most canonical PBMC populations (Fig. 2a, upper right), yet substantial overlap persists among closely related immune states, including naïve and memory CD4⁺ T cells, CD4⁺ and CD8⁺ T cells, and monocyte subtypes. These regions of overlap reflect intrinsic uncertainty at the single-cell resolution and illustrate a key limitation of per-cell inference: local evidence alone is insufficient to enforce coherent population structure.

To incorporate population structure, MOSAIC aggregates cell-level probability vectors within Leiden clusters^24^ to obtain cluster-wise soft consensus profiles. Assigning clusters according to their dominant consensus probability markedly sharpens the separation between major immune lineages and suppresses scattered misassignments (Fig. 2a, lower left). Importantly, this stabilization arises from aggregating full probability distributions rather than hard labels, allowing consistent but weak lineage signals to accumulate while preserving information about competing alternatives.

Beyond consensus assignment, MOSAIC applies an adaptive decision module that evaluates evidence strength and ambiguity at the cluster level. This module jointly considers the magnitude of the top-ranked lineage probability, separation from competing candidates, and relative enrichment across clusters. Based on these global criteria, clusters are classified as Confident, Mixed (A/B), or Unknown. In the adaptive representation, confidently resolved clusters retain distinct lineage identities, whereas Mixed and Unknown clusters are collapsed into a neutral background (Fig. 2a, lower middle), explicitly marking regions where marker evidence is ambiguous or insufficient and where the framework deliberately avoids overconfident labeling.

Cluster-level decisions are then reconciled with local cell-level evidence to produce refined cell-level annotations. Only cells that are consistent with their cluster identity and exhibit sufficient individual support are assigned a final label, while cells from uncertain clusters or with weak evidence remain marked as ambiguous (Fig. 2a, lower right). This refinement enforces coherence between local heterogeneity and global organization while preserving uncertainty where warranted.

The biological validity of these adaptive labels is supported by canonical marker expression patterns. UMAP overlays of representative lineage markers, including CCR7, CD8A, CD14, and GZMB, display spatially coherent and lineage-specific expression domains that align closely with the adaptively inferred cluster identities (Fig. 2b). CCR7 marks naïve and memory CD4⁺ T-cell populations, CD8A highlights cytotoxic lineages, CD14 identifies classical monocytes, and GZMB localizes to effector CD8⁺ T cells and natural killer cells, collectively recapitulating well-established immune organization.

Consistently, dot-plot summaries aggregated by adaptive cluster labels reveal distinct and biologically expected marker signatures across immune compartments (Fig. 2c). Both marker intensity and the fraction of expressing cells confirm that each adaptively defined cluster exhibits a coherent transcriptional profile, indicating that the inferred identities are marker-supported rather than algorithmically imposed.

Together, this PBMC example demonstrates that it is the structured integration of probabilistic cell-level evidence, cluster-level consensus, and adaptive uncertainty-aware decisions—rather than any single component in isolation—that enables MOSAIC to generate stable, interpretable, and biologically grounded annotations. This multi-level design forms the conceptual foundation for the accuracy, robustness, and uncertainty characterization explored in subsequent analyses.

To assess whether the proposed multi-level inference structure extends beyond immune-specific cellular hierarchies, we applied MOSAIC to liver tissue as a representative non-immune system. In this setting, cell-level probability profiles preserve local ambiguity, while robust and coherent population-level annotations emerge through cluster-level integration, consistent with the framework’s underlying design principles (Extended Data Fig. 1).

### Result 2 | Multi-level probabilistic aggregation improves annotation accuracy across tissues and paradigms

To evaluate the practical consequences of multi-level probabilistic integration, we benchmarked MOSAIC against representative annotation methods spanning major methodological paradigms: the reference-based approach SingleR¹³, pretrained classifiers CellTypist¹⁵ and scDeepSort¹⁶, and marker-rule–based systems scCATCH²⁵ and scType¹¹. Benchmarking was performed across six real tissues—PBMC, liver, colon, pancreas, lung, and retina—using consistent evaluation protocols and default parameter settings.

At the population level, MOSAIC achieved the highest or near-highest annotation accuracy across all tissues (Fig. 3a, top panels). Performance gains are most pronounced in datasets characterized by closely related or partially overlapping cell states, such as PBMC and lung. In these complex immune environments, adaptive cluster-level inference preserved fine-grained separation among T-cell and monocyte subsets more effectively than competing approaches, indicating improved resolution in heterogeneous cell populations.

**Figure 3.**
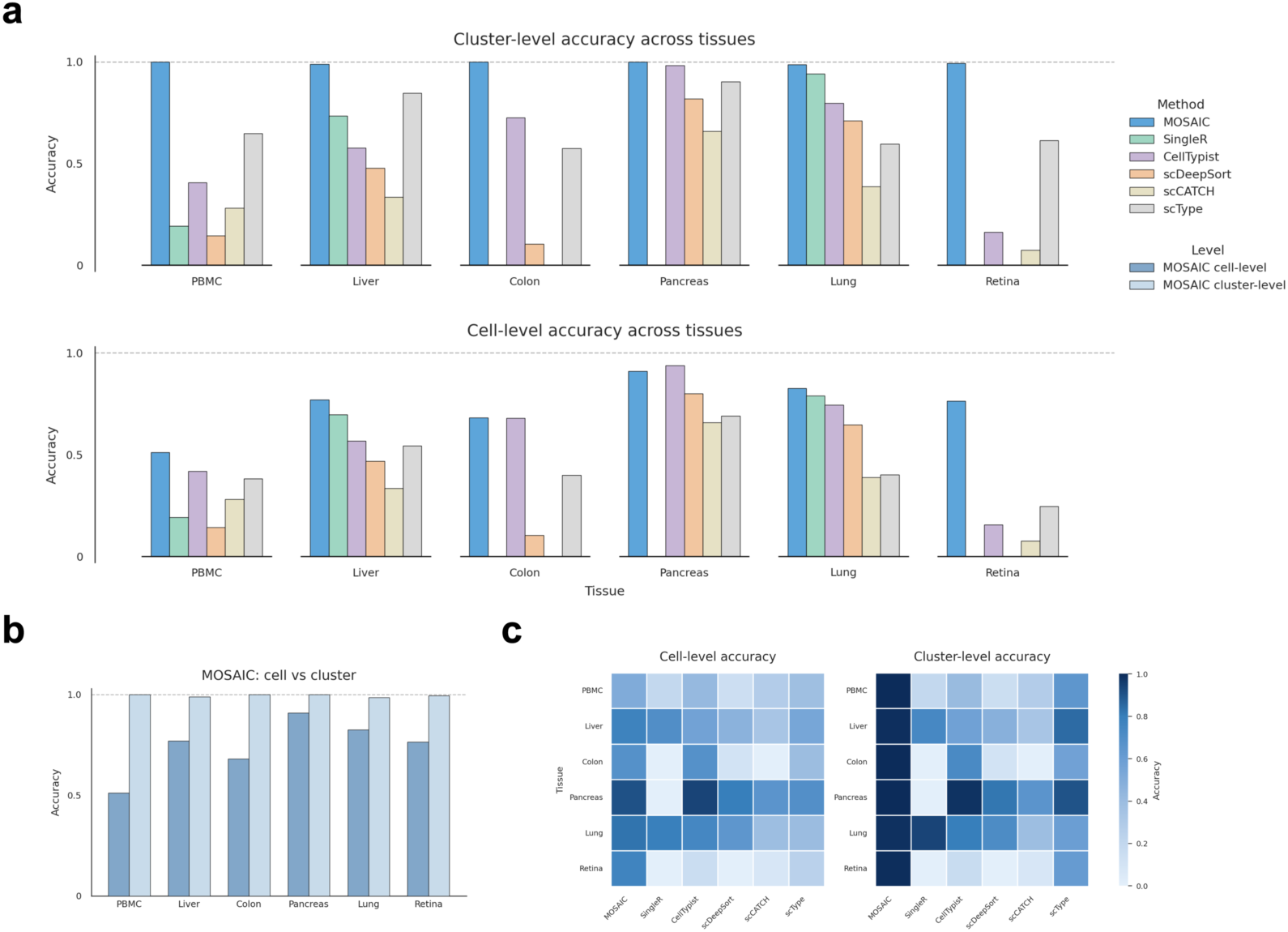
Population-level probabilistic aggregation improves annotation stability and accuracy across datasets. **(a)** Cluster-level (top) and cell-level (bottom) annotation accuracy across six real tissues (PBMC, liver, colon, pancreas, lung, and retina). MOSAIC is compared with representative reference-based (SingleR), large-scale classifier (CellTypist), deep-learning (scDeepSort), and marker-rule–based (scCATCH, scType) methods under matched annotation granularity. Cluster-level accuracy corresponds to labels derived from population-level probabilistic consensus, whereas cell-level accuracy corresponds to maximum a posteriori (argmax) decoding of cell-level probability vectors. **(b)** Direct comparison of MOSAIC cell-level argmax predictions and cluster-level adaptive annotations across tissues. Cell-level predictions exhibit substantial variability and reduced accuracy across organs, whereas cluster-level inference consistently achieves higher and more stable accuracy in all six tissues, demonstrating that population-level probabilistic aggregation is a necessary component for coherent annotation beyond independent per-cell classification. **(c)** Heatmap summary of cell-level (left) and cluster-level (right) accuracy for all methods and tissues, illustrating cross-tissue performance patterns and differences among annotation paradigms. Additional performance metrics, including Macro-F1, ARI, and NMI, are provided in Supplementary Tables S1–S4.

At the cell level, argmax predictions derived from probabilistic scoring already provided competitive accuracy relative to external tools in most tissues (Fig. 3a, bottom panels). However, a direct comparison between cell-level argmax predictions and cluster-level adaptive annotations reveals a systematic and consistent pattern: cell-level predictions alone are less stable and less accurate, whereas cluster-level probabilistic aggregation uniformly improves performance across all datasets (Fig. 3b). In every tissue examined, cluster-level accuracy exceeds cell-level accuracy, demonstrating that population-level integration is not a cosmetic refinement but a necessary component for coherent and reliable annotation. This improvement reflects the statistical effect of aggregating full probability distributions rather than hard labels. By pooling probabilistic marker evidence across cells, MOSAIC suppresses scattered single-cell misassignments while amplifying consistent lineage signals, thereby stabilizing inference without sacrificing sensitivity to biologically meaningful heterogeneity.

Heatmap summaries of accuracy (Fig. 3c) further reveal characteristic differences among annotation paradigms. Reference-based and pretrained classifier methods exhibit pronounced tissue dependence, reflecting sensitivity to domain mismatch and training data bias. Marker-rule–based approaches are constrained by predefined marker sets and often underperform in heterogeneous tissues with overlapping identities. In contrast, MOSAIC maintains uniformly high performance at both cell and cluster levels across all organs, reflecting the benefit of integrating probabilistic cell-level evidence with population-level structure rather than relying on single-resolution decision rules. All quantitative benchmarking results were obtained using default parameter settings across tissues, with comprehensive performance summaries provided in Supplementary Tables S1– S4.

Together, these results show that improved annotation accuracy emerges as a direct consequence of structured multi-level probabilistic aggregation. Rather than relying on increasingly complex cell-level classifiers, MOSAIC demonstrates that enforcing coherence through population-level inference provides a more stable and generalizable solution across heterogeneous datasets and annotation paradigms.

### Result 3 | Probabilistic uncertainty reveals transitional states and enables principled Mixed and Unknown assignments

Most existing cell-type annotation methods return deterministic labels without an explicit representation of uncertainty, limiting their ability to quantify ambiguity or represent graded transitions, even in well-characterized datasets such as PBMC (Extended Data Fig. 2). In such settings, uncertainty is typically inferred indirectly from label disagreement or ad hoc confidence scores, rather than modeled as an intrinsic component of the annotation process. In contrast, MOSAIC retains the full posterior probability vector for every cell, enabling uncertainty to be quantified directly within the model. From these probability distributions, we derived complementary uncertainty measures, including normalized entropy, maximum class probability, and the separation between the top-ranked and second-ranked candidates.

When projected onto the UMAP embedding, these metrics exhibit a coherent spatial organization: cells with elevated uncertainty consistently localize to boundary regions between closely related immune states, including the interfaces between naïve and memory CD4⁺ T cells, between memory CD4⁺ and effector CD8⁺ T cells, and between classical and non-classical monocytes (Fig. 4a). This structured pattern indicates that uncertainty is not randomly distributed but instead reflects biologically structured intermediate states aligned with known immune differentiation continua^6,7,8^.

**Figure 4.**
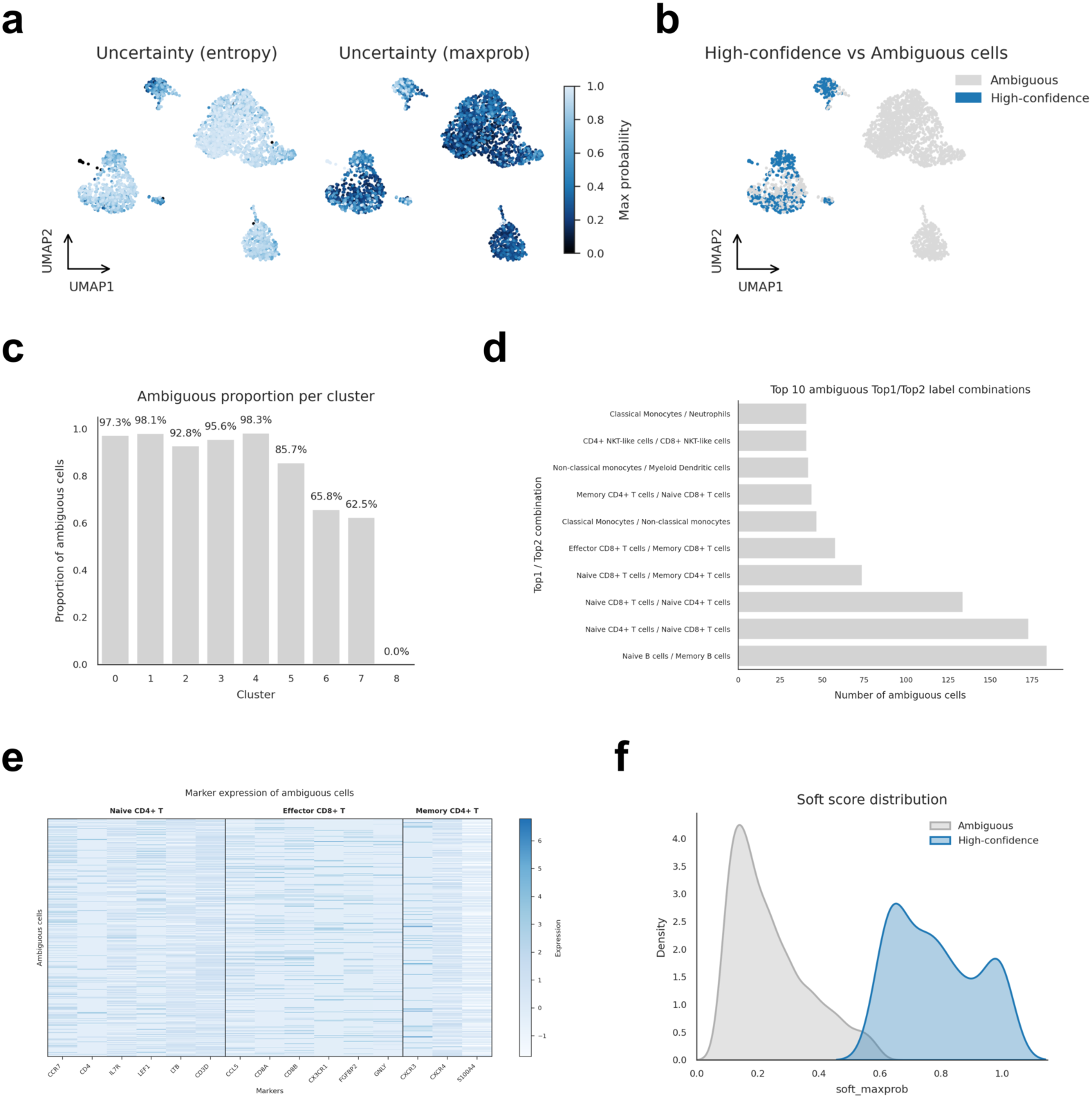
Probabilistic uncertainty is structured and enables principled identification of ambiguous and transitional states. **(a)** UMAP projections of cell-level uncertainty measures derived from full posterior probability vectors, including normalized entropy (left) and maximum class probability (right). Elevated uncertainty consistently localizes to boundary regions between closely related immune states, whereas transcriptionally well-separated populations exhibit low entropy and high maximum class probability. **(b)** UMAP visualization of high-confidence versus ambiguous cells defined by a global threshold on probability strength. Ambiguous cells concentrate at lineage boundaries highlighted by entropy and maximum class probability maps, while high-confidence cells occupy transcriptionally well-separated regions. **(c)** Fraction of ambiguous cells within each Leiden cluster, revealing substantial heterogeneity across clusters. Ambiguity is enriched in clusters adjacent to closely related identities and depleted in transcriptionally well-resolved populations, indicating population-level structure in uncertainty distribution. **(d)** Top ten most frequent top1–top2 label combinations among ambiguous cells. Dominant ambiguous pairings occur between biologically related immune identities, including naïve versus memory CD4⁺ T cells, memory CD4⁺ versus effector CD8⁺ T cells, and classical versus non-classical monocytes, demonstrating that uncertainty is structured along meaningful lineage axes rather than arising from random misclassification. **(e)** Heatmap of marker expression profiles for ambiguous cells across representative immune lineages. Ambiguous cells exhibit mixed expression of canonical markers associated with neighboring cell states, consistent with intermediate or transitional transcriptional programs that cannot be faithfully captured by discrete labels. **(f)** Distribution of maximum class probability for ambiguous and high-confidence cells. Ambiguous cells display a pronounced shift toward lower maximum probability values, whereas high-confidence cells cluster near unity, indicating that uncertainty is probabilistically well resolved within the model rather than reflecting stochastic variation. For PBMC visualization only, a more permissive probability threshold was applied to improve interpretability of ambiguous cells; visualization parameters are identical to those used in Fig. 2. This threshold was used exclusively for visualization and did not affect uncertainty quantification, cluster classification, or any downstream analysis.

Maximum class probability maps further illustrate this distinction. Cells within well-separated clusters display uniformly high confidence, whereas cells positioned at lineage boundaries show markedly reduced maximum class probability (Fig. 4a). Applying global probability thresholds allows cells to be classified as high-confidence or ambiguous without reference to external calibration models. When visualized on the UMAP, ambiguous cells concentrate precisely at the same transitional regions highlighted by entropy-based measures (Fig. 4b), demonstrating that uncertainty captured by MOSAIC reflects intrinsic structure in the data rather than technical noise or stochastic misclassification.

The distribution of ambiguous cells across clusters provides additional support for this interpretation. Some clusters contain disproportionately high fractions of ambiguous cells, whereas others are almost entirely composed of high-confidence assignments (Fig. 4c). This heterogeneity reflects differences in population composition: ambiguity is enriched in clusters adjacent to closely related lineages and depleted in transcriptionally well-separated populations.

Importantly, uncertainty is not arbitrary but structured along biologically meaningful axes. Examining the dominant top1–top2 label combinations among ambiguous cells reveals that the most frequent ambiguous pairings occur between neighboring immune identities, including naïve versus memory CD4⁺ T cells, memory CD4⁺ versus effector CD8⁺ T cells, and classical versus non-classical monocytes (Fig. 4d). These pairings correspond closely to established immune maturation and activation trajectories, reinforcing the interpretation that ambiguous assignments reflect intermediate phenotypes rather than annotation error^6,7,23^.

At the transcriptional level, ambiguous cells exhibit mixed marker expression patterns that bridge canonical lineages (Fig. 4e). Cells ambiguous between T-cell subtypes co-express markers associated with both naïve and effector programs, including CCR7, IL7R, CCL5, GZMB, CXCR3, and S100A4. This mixed expression profile supports the view that these cells represent activation-associated or differentiation-intermediate states that cannot be faithfully captured by discrete categorical labels^8,23^.

Finally, probability distributions provide a clear quantitative separation between ambiguous and high-confidence populations. Ambiguous cells exhibit a pronounced left shift in maximum class probability, whereas high-confidence cells cluster near unity (Fig. 4f). This separation demonstrates that uncertainty is probabilistically well resolved within the model, rather than arising from arbitrary thresholding or stochastic variation. Calibration analyses^26^ show that annotation accuracy varies systematically with binned maximum class probability across tissues, indicating that probabilistic scores produced by MOSAIC capture meaningful confidence information rather than arbitrary values (Supplementary Fig. S1).

Together, these results highlight a fundamental distinction between collapsing intermediate biological states into fixed categorical labels and explicitly modeling uncertainty through probabilistic representations. By treating uncertainty as a first-class component of annotation, MOSAIC enables principled identification of confident, ambiguous, and transitional cellular states within a unified inference framework.

Notably, similarly structured patterns of cell-level uncertainty are observed in liver tissue (Extended Data Fig. 3), where elevated uncertainty consistently localizes to interfaces between transcriptionally related populations and mixed cellular compartments. This spatial organization mirrors that observed in PBMC and indicates that probabilistic ambiguity in MOSAIC arises from structured evidence integration rather than from immune-specific biology. Consistent uncertainty patterns are further observed across additional non-immune tissues, including lung, pancreas, colon, and retina (Supplementary Fig. S2), supporting the generality of the underlying inference behavior across diverse tissue contexts.

Together, these analyses demonstrate that uncertainty in MOSAIC is structured, reproducible, and grounded in marker evidence across tissues. Rather than treating ambiguity as noise or classification failure, MOSAIC explicitly models uncertainty as an intrinsic component of the annotation process, enabling gradual transitions and mixed identities to be represented directly within the probabilistic framework. Importantly, Mixed and Unknown states emerge naturally from the multi-level inference design, without reliance on ad hoc post-processing or external calibration, and therefore correspond to coherent intermediate or transitional cellular states rather than methodological artifacts.

### Result 4 | Controlled dropout perturbations validate the robustness of multi-level probabilistic inference

To systematically evaluate robustness under progressive signal degradation, we conducted controlled dropout perturbation experiments using scDesign2^27^. Increasing levels of expression dropout (0–0.8) were introduced across six tissues (PBMC, lung, liver, colon, pancreas, and retina), gradually suppressing marker signal while preserving global population structure. These perturbations provide a stringent and biologically grounded test of whether multi-level probabilistic integration remains valid when cell-level evidence becomes increasingly sparse^28^.

Across all tissues and perturbation levels, MOSAIC maintains consistently high annotation performance (Fig. 5a). In PBMC, lung, liver, and colon—datasets characterized by closely related cell states and substantial transcriptional overlap—accuracy remained remarkably stable under mild to moderate dropout (0–0.6), followed by gradual and structured declines as signal loss increased further. Even at the highest perturbation level (0.8), MOSAIC consistently ranked among the best-performing methods across tissues. In contrast, several competing approaches exhibit sharper performance collapse and increased instability in heterogeneous settings. Pancreas, which is dominated by transcriptionally well-separated endocrine populations, showed uniformly high accuracy across multiple methods, serving as a control regime in which fine-grained inference is less demanding.

**Figure 5.**
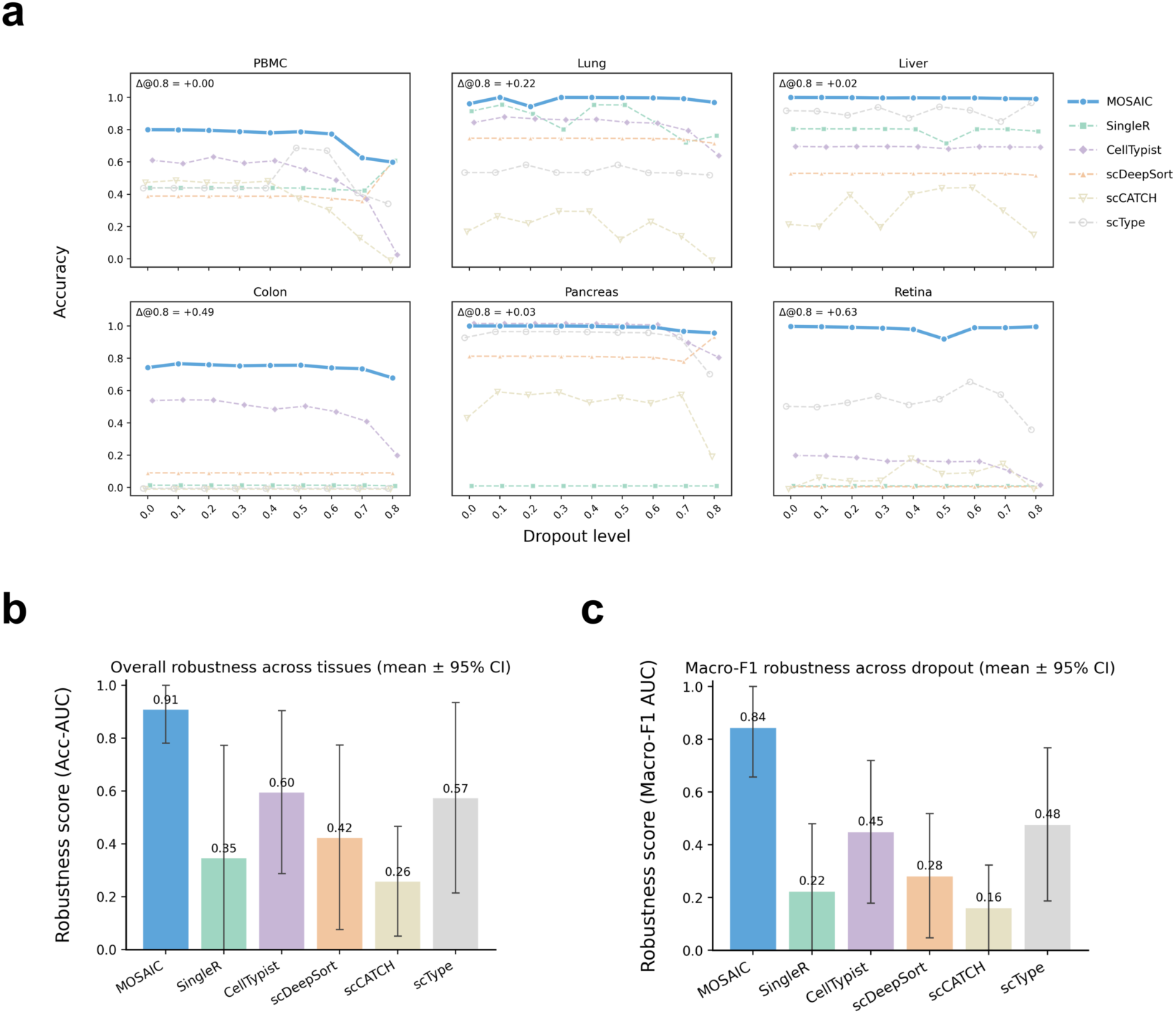
Controlled dropout perturbations validate the robustness of multi-level probabilistic inference. **(a)** Annotation accuracy as a function of increasing scDesign2-simulated dropout levels (0–0.8) across six tissues (PBMC, lung, liver, colon, pancreas, and retina). Progressive dropout suppresses observed gene expression by introducing additional sparsity, MOSAIC maintains high accuracy across tissues, whereas competing methods exhibit sharper performance collapse or persistently lower accuracy, particularly in heterogeneous datasets. Pancreas, dominated by transcriptionally well-separated endocrine populations, serves as a control regime in which most methods achieve high accuracy. Δ@0.8 denotes the absolute accuracy difference between MOSAIC and the best-performing competing method at the highest dropout level (0.8) for each tissue. **(b)** Overall robustness quantified as the area under the accuracy–dropout curve (Acc-AUC), averaged across tissues. Bars indicate mean robustness scores, and error bars represent t-distribution–based 95% confidence intervals computed across tissues and clipped to the valid range [0,1] for visualization, reflecting the bounded nature of AUC-based robustness metrics. MOSAIC achieves the highest Acc-AUC among all evaluated methods, indicating sustained annotation performance under progressive marker loss. **(c)** Robustness of cell-type resolution assessed using the area under the Macro-F1–dropout curve (Macro-F1 AUC). Values represent tissue-averaged means with t-distribution–based 95% confidence intervals, clipped to [0,1] for visualization. MOSAIC consistently outperforms reference-based, classifier-based, deep-learning, and marker rule–based approaches, demonstrating that robustness is preserved across diverse cell types rather than being driven by dominant populations. Together, these analyses show that robustness in MOSAIC emerges from structured probabilistic aggregation and adaptive population-level inference, which preserve discriminative structure as marker evidence is progressively degraded.

Importantly, robustness is not fully characterized by absolute accuracy alone, but by how performance degrades as marker signal is progressively suppressed. Several baseline methods exhibit relatively flat but low accuracy plateaus across dropout levels. Under a comparable annotation granularity, this behavior reflects persistent misclassification rather than genuine robustness: as marker information is degraded, these methods lose discriminative power but continue to produce deterministic assignments, leading to stable yet systematically incorrect predictions. In contrast, MOSAIC maintains a substantially higher accuracy plateau across perturbation levels, indicating that probabilistic evidence integration and population-level aggregation preserve discriminative structure under signal loss, rather than merely freezing early errors.

To summarize robustness across tissues, we quantified the area under the accuracy–dropout curves and report mean robustness scores with 95% confidence intervals (Fig. 5b). MOSAIC achieved the highest overall robustness score among all evaluated methods, demonstrating sustained stability across diverse tissue contexts. This advantage was further supported by tissue-averaged Macro-F1 robustness across dropout levels (Fig. 5c), indicating that performance gains are maintained across cell types rather than being driven by dominant populations alone.

These results indicate that robustness in MOSAIC emerges from its structured multi-level inference design. Probabilistic cell-level representations preserve uncertainty as marker evidence degrades^5,29^ and serve as calibrated inputs for population-level aggregation. Cluster-level soft consensus stabilizes inference by amplifying consistent signals across cells, while adaptive refinement prevents overconfident assignments when evidence becomes insufficient. Together, these components enable MOSAIC to maintain coherent and biologically meaningful annotations under severe perturbation, validating the framework’s suitability for noisy, low-depth, or technically challenging single-cell datasets. Additional robustness analyses and tissue-specific results are provided in Supplementary Figs. S3–S8 and Supplementary Tables S5–S9.

## Discussion

This study presents MOSAIC as a structured multi-level framework that reframes cell-type annotation as a coordinated inference problem rather than a collection of independent classification decisions. Most current strategies, whether marker-based, reference-based, or machine-learning–based, operate at a single analytical resolution and struggle to reconcile fine-grained single-cell variation with higher-order population structure^1,2,10,30^. As a result, these methods tend to produce deterministic labels that obscure uncertainty, fail to represent gradual or mixed identities, and lack coherence across biological scales^5,7,9,31^. MOSAIC resolves this disconnect by explicitly integrating marker evidence, probabilistic reasoning, and cluster-level organization within a unified multi-level framework, consistent with growing evidence that population context and multi-scale structure are essential for robust single-cell inference^12,17,21,22^. From a methodological standpoint, this work directly motivates the necessity of a cluster-level layer: without such integration, cell-level predictions remain sensitive to uncertainty and insufficient to support coherent biological annotation^12,20^.

A first conceptual contribution of MOSAIC is the formulation of a two-layer, direction-aware marker scoring scheme that generalizes classical marker-based annotation into a structured inference setting. Drawing on established marker-based principles, particularly the joint use of positive and negative markers formalized by scType^11^, MOSAIC extends this cell-level paradigm into a structured multi-level system. By jointly modeling positive and negative markers, the framework extends traditional marker interpretation beyond categorical cell-level decisions to a coherent multi-resolution process. At the single-cell level, direction-aware scoring yields quantitative enrichment profiles that preserve both lineage-specific evidence and uncertainty. At the population level, aggregation of these profiles stabilizes inference by amplifying consistent but weak signals that are often masked by dropout and stochastic variation. This explicit separation of evidence generation and evidence integration restores biological separability in systems characterized by gradual differentiation or substantial technical noise, and provides a principled substrate for downstream probabilistic reasoning.

A second major contribution is the use of dual-layer probabilistic soft labeling as a central representational choice. Retaining full probability vectors at both the cell and cluster levels allows uncertainty to be quantified, propagated, visualized, and interpreted directly, aligning with foundational perspectives on uncertainty as an intrinsic component of predictive modeling rather than a nuisance parameter^5,32,33^. At the single-cell level, probabilistic outputs capture low-confidence and intermediate states that deterministic classifiers tend to collapse, consistent with observations from trajectory and fate-probability analyses showing that cell identity often lies on a continuum^6,7^. At the cluster level, soft consensus profiles distinguish well-resolved populations from those containing competing lineage signals. Together, these representations elevate uncertainty to a structured and biologically meaningful property of the data, rather than as noise or classification failure. Crucially, this uncertainty often corresponds to stable intermediate or transitional cellular states that arise from continuous biological processes, particularly evident in immune systems but not restricted to them. ^8,23,34^.

The third contribution of MOSAIC lies in its multi-level adaptive refinement strategy, which translates probabilistic representations into interpretable annotation outcomes. By jointly considering marker strength, probability separation, relative enrichment, and cross-level consistency, the framework assigns clusters to Confident, Mixed, or Unknown categories and propagates these decisions back to individual cells. This strategy mitigates long-standing challenges in annotating ambiguous populations, including Mono–DC, T–NK, ductal–acinar, and rod–cone intermediates, and avoids overconfident assignments when evidence is limited^35,36,37^. As a result, refined labels preserve coherence between local single-cell variation and global population structure, yielding annotations that align with known biological continua.

Across six real tissues and controlled scDesign2 dropout perturbations, MOSAIC consistently matches or outperforms representative methods spanning all major annotation paradigms, including reference-based^13^, machine-learning^15^, deep-learning^16^, and marker-rule systems^25^. Importantly, these improvements are not limited to classification accuracy.

MOSAIC uniquely integrates interpretability, robustness to noise, explicit uncertainty modeling, and cross-level consistency—capabilities that existing approaches typically address in isolation, if at all^38,39,40,41^. The resulting annotations exhibit reduced scattered errors, improved population coherence, sharper population boundaries, and a more faithful representation of transitional or mixed cellular states.

In summary, this work advances cell-type annotation beyond single-level prediction by explicitly modeling how probabilistic cell-level signals aggregate into population-level structure, as instantiated through cluster-level organization, and how that structure, in turn, refines cell-level decisions. By formulating cell-type annotation as a structured multi-level inference problem rather than a single-resolution classification task, MOSAIC provides a generalizable computational framework for interpreting complex and heterogeneous single-cell datasets. This perspective resolves fundamental limitations of existing approaches and establishes uncertainty-aware, interpretable annotation as a first-class modeling objective in large-scale single-cell analysis.

## Methods

### Method 1 | Dataset processing and preprocessing

Six publicly available single-cell RNA sequencing (scRNA-seq) datasets representing diverse tissues and cellular compositions were analyzed in this study, including peripheral blood mononuclear cells (PBMC; 10x Genomics PBMC 3k dataset, as used in the Seurat guided clustering tutorial)^42^, lung (Weizmann Institute 3CA lung atlas; Qian et al., 2020)^43^, retina (GSE137537), colon (GSE81547), pancreas (GSE116222), and liver (GSE124395). These datasets encompass both immune- and parenchymal-dominated tissues and span a range of transcriptional heterogeneity, providing a diverse setting for evaluating the behavior of multi-level probabilistic annotation under heterogeneous biological contexts. All datasets were processed using a unified preprocessing pipeline implemented^44^ in Scanpy (v1.9.8) to ensure consistency across tissues.

For each dataset, cells with fewer than 200 detected genes were removed to exclude low-quality or empty droplets, and genes expressed in fewer than three cells were discarded. Mitochondrial genes were identified based on the “MT-” prefix, and standard quality-control metrics—including total counts, number of detected genes, and mitochondrial gene fraction—were computed for diagnostic purposes. Unless otherwise stated, no dataset-specific filtering beyond these criteria was applied.

Raw count matrices were normalized to a total of 10,000 counts per cell and log-transformed using a natural logarithm. Highly variable genes were selected using the Scanpy implementation of the Seurat v3 variance-stabilizing method^17^, retaining 2,000 genes per dataset. Gene expression values were subsequently scaled to unit variance without clipping.

Dimensionality reduction and neighborhood graph construction followed standard Scanpy workflows. Principal component analysis (PCA) was performed using 10–30 components, depending on dataset size and complexity. A k-nearest-neighbor graph was constructed in PCA space, and Leiden clustering was applied with dataset-specific resolution parameters (0.2–0.6) chosen to capture major biological compartments without inducing excessive cluster fragmentation^24^. Uniform Manifold Approximation and Projection (UMAP) embeddings were generated solely for visualization and did not influence downstream scoring or annotation steps^45^.

### Method 2 | Marker databases and gene-set construction

Marker-based annotation in this study builds on established principles of lineage-specific gene expression while extending them into a multi-level probabilistic framework. For all six tissues analyzed, we assembled a fixed set of curated marker genes used to score annotated cell types, with explicit separation of positive and negative markers.

Initial marker lists were derived from a combination of widely used community resources, including CellMarker^46^, PanglaoDB^47^, and the Human Protein Atlas^48^, together with markers reported in primary literature and domain knowledge for immune, epithelial, endothelial, and stromal lineages. In addition, we adopted the conceptual distinction between positive and negative markers formalized by scType^11^, which demonstrated that incorporating mutually exclusive lineage markers improves interpretability at the single-cell level. Gene symbols were standardized to HGNC conventions, and tissue-specific aliases were unified to ensure consistent mapping across datasets.

Following established marker-based annotation principles, most notably those formalized by scType, each annotated cell type was represented by complementary gene sets capturing positive lineage evidence and, where informative, negative lineage exclusion:

i. positive markers, consisting of genes preferentially enriched in the focal lineage; and
ii. negative markers, consisting of genes characteristic of competing or mutually exclusive lineages and not expected to be expressed in the target cell type.

Unlike traditional marker-based schemes that rely solely on positive evidence, this dual-direction design enables direction-aware scoring by simultaneously rewarding lineage-consistent expression and penalizing conflicting transcriptional signatures. Importantly, marker selection was performed at the level of broad biological lineages rather than optimized for individual datasets, thereby avoiding dataset-specific tuning.

Marker lists were further refined through manual curation to remove genes with ubiquitous expression, strong cell-cycle or stress associations, or inconsistent annotation across studies, following general principles for robust marker selection and evaluation^49^. This process yielded compact, interpretable marker sets that emphasize lineage identity rather than dataset-dependent expression artifacts. The final marker dictionaries were indexed by tissue and cell type and applied uniformly across all real datasets, benchmarking analyses, and dropout simulation experiments.

To ensure fair comparison with existing annotation methods, marker sets were fixed a priori and were not modified based on downstream performance. All competing methods were evaluated using their recommended default configurations and reference resources. Performance gains observed for MOSAIC therefore arise from the structured multi-level integration of marker evidence and probabilistic population-level reasoning, rather than from expanded or task-specific marker definitions.

### Method 3 | Multi-level marker scoring and probabilistic representation

Our framework performs marker-based annotation through a structured, multi-level scoring scheme that explicitly separates cell-level evidence extraction from population-level signal stabilization^12^. Direction-aware marker scoring is first applied to individual cells to quantify lineage-specific support and conflict, after which probabilistic evidence is aggregated within clusters to derive low-variance population-level summaries. This design enables uncertainty-aware inference while preserving biological interpretability across scales^5,33^.

#### Cell-level direction-aware marker scoring

For each cell *i* and candidate cell type *t*, we define a set of positive markers *M_t_*^+^ and, when available, a set of negative markers *M*_t_^−^, Positive markers correspond to genes preferentially enriched in the focal lineage, whereas negative markers represent genes characteristic of competing or mutually exclusive cell types and are not expected to be expressed in the target lineage. This formulation follows established marker-based annotation principles, most notably those formalized by scType, but is generalized here into a probabilistic framework.

Let *x*_*ig*_ denote the log-normalized expression of gene *g* in cell *i*. The positive-marker score is computed as a size-normalized aggregate of expression across the positive marker set, which preserves cumulative support from multiple markers while preventing linear inflation with marker set size:

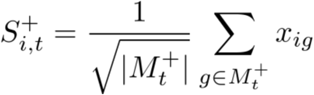

Similarly, when negative markers are defined, a negative-marker score is computed as a uniform exclusion term:

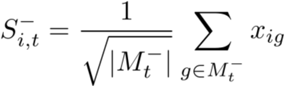

Positive and negative evidence are then combined into a cell-level logit:

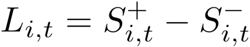

This direction-aware formulation jointly captures supportive and conflicting marker evidence while reducing sensitivity to stochastic dropout by aggregating signals across gene sets rather than relying on individual markers.

To obtain probabilistic outputs, logits are transformed using a temperature-controlled softmax:

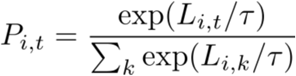

where *τ* controls distribution sharpness (default *τ* =1).

The resulting probability matrix *P*∈ℝ^*N*×*T*^, where *N* denotes the number of cells and *T* the number of candidate cell types, provides a full probabilistic characterization of lineage support for each cell, retaining uncertainty information that is lost in deterministic label assignment^5,32,33,50,51^.

#### Cluster-level probability aggregation

To incorporate population-level structure and improve robustness to single-cell noise, cell-level probability vectors are aggregated within each Leiden cluster *C*_*k*_. For each cluster *k* and candidate cell type *t*, a cluster-level consensus probability is computed as:

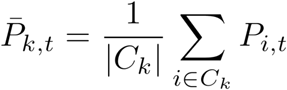

This aggregation produces a low-variance summary of marker support that stabilizes lineage signals under technical noise and dropout, while preserving information about competing identities within the cluster. Unlike majority voting over hard labels, aggregation operates directly on full probability vectors, allowing dominant and secondary lineage signals to be retained.

The top-ranked type

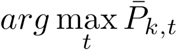

is stored as the cluster-level soft consensus label, while the full consensus vector *P̄*_*k*_ is retained for downstream uncertainty-aware decision-making.

These cluster-level probabilistic summaries constitute the stabilized population-level evidence used by the adaptive refinement module described below^52^. Importantly, all downstream uncertainty quantification, adaptive cluster-level decisions, and refined cell-level annotations are derived exclusively from these probabilistic representations, without introducing additional scoring heuristics or post hoc corrections.

### Method 4 | Adaptive cluster-level decision module

To translate cluster-level probabilistic summaries into interpretable annotation outcomes, MOSAIC employs an adaptive decision module that assigns each Leiden cluster^24^ to one of three categories—Confident, Mixed, or Unknown— based on complementary statistical properties of aggregated marker evidence^35,36,37,53^.

Notably, adaptive cluster labels are derived directly from aggregated direction-aware marker scores rather than from probability vectors, ensuring that population-level decisions reflect the same underlying scoring mechanism used for cell-level evidence extraction.

For each Leiden cluster *C*_*k*_, direction-aware marker scores *L*_*i*,*t*_ for cells *i* ∈ *C*_*k*_ are aggregated to quantify cluster-level support for each candidate cell type. Specifically, aggregated support is computed as the sum of marker scores across cells and normalized by the number of constituent cells, yielding a per-cell support estimate:

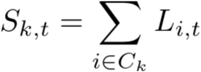

This per-cell normalized aggregation ensures comparability across clusters of different sizes while preserving the cumulative strength of lineage-consistent marker evidence.

Three complementary statistics are computed from these cluster-level summaries to jointly characterize the strength, separability, and specificity of marker support. Together, these criteria prevent confident cluster assignments driven solely by strong but non-specific signals or by marginal separation in the absence of sufficient evidence.

First, the overall strength of support for the leading candidate identity is quantified as

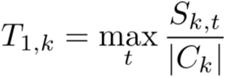

which reflects the average level of lineage-consistent marker evidence accumulated across cells within the cluster.

Second, to assess whether the leading identity is clearly separated from alternative candidates, the margin between the highest- and second-highest supported identities is computed. Let *t_1_* and *t_2_* denote these two candidates, respectively:

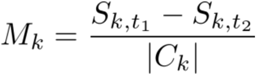

This margin captures the degree of internal ambiguity within the cluster, distinguishing dominant lineages from competing or transitional states^37^.

Third, to evaluate whether the leading identity is specifically enriched in the focal cluster relative to the global population, a relative enrichment statistic is computed as

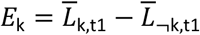

where *L̄*_k,t–1_ and *L̄*_*k*,*t*–_*_1_* denote the mean marker scores for type *t_1_* inside and outside the cluster, respectively.

Rather than applying fixed thresholds, each statistic is compared against dataset-specific quantile thresholds computed across all clusters within the same dataset. Specifically, clusters with overall support *T_1_*_,*k*_ falling below the lower quantile are assigned Unknown labels, reflecting insufficient or inconsistent marker evidence. Among remaining clusters, those with inadequate separation *M*_*k*_ are classified as Mixed, indicating competing lineage support. Clusters exhibiting strong support, clear separation, and sufficient enrichment are labeled Confident with a single dominant identity.

These adaptive cluster-level labels provide a population-level context for downstream refinement rather than final cell-level assignments. Sensitivity of adaptive labeling to Leiden resolution^24,54^ was evaluated separately and is reported in Supplementary Table S10, demonstrating stable performance across a broad range of clustering granularities.

### Method 5 | Cell-level refinement module

The final output of MOSAIC is a refined cell-level annotation that explicitly distinguishes Confident cell assignments from Ambiguous ones. Importantly, this refinement step is not a standalone confidence filter applied to cell-level predictions, but a cluster-conditioned inference procedure that integrates cell-level probabilistic evidence with adaptive cluster-level decisions derived in the previous step^35,36,37,51^.

Let *P*_*i*_ denote the cell-level probability vector for cell i, obtained from the temperature-controlled softmax transformation of direction-aware marker scores (Section 3), and let *C*_*k*_ denote the Leiden cluster to which cell *i* belongs. Let *L*_*k*_∈ {Confident, Mixed, Unknown} denote the adaptive label assigned to cluster *C*_*k*_ by the cluster-level decision module (Section 4).

#### Cells in Confident clusters

For clusters labeled Confident, a single dominant lineage *t* has been identified at the population level based on strong marker support, clear separability, and cluster-specific enrichment. In this setting, MOSAIC permits cell-level commitment only when individual evidence is both strong and consistent with the cluster-level identity.

Specifically, a cell *i* belonging to a Confident cluster with dominant lineage *t* is assigned a Confident cell-level label *t* if and only if:

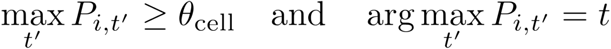

where *θ*_*cell*_ is a fixed probability threshold controlling the stringency of cell-level commitment.

Unless otherwise stated, a fixed threshold (*θ*_*cell*_ = 0.8) was applied uniformly across tissues and experiments to ensure comparability. Crucially, this threshold does not influence probability estimation and is applied only after probabilistic inference, serving exclusively as a commitment criterion for translating probabilistic evidence into final labels.

Importantly, *θ*_*cell*_ is not a structural assumption of the framework but an implementation-level parameter that controls the stringency of cell-level commitment within a cluster-conditioned inference scheme. Lower values allow more permissive resolution of individual cells, whereas higher values enforce stricter consistency with cluster-level context.

Consistent with its intended role as a commitment parameter rather than a driver of probabilistic scoring, moderate variations of *θ*_*cell*_ primarily affected the fraction of cells labeled Confident, while overall annotation accuracy and clustering agreement metrics remained stable (Supplementary Table S11).

Cells that do not satisfy both conditions are labeled Ambiguous, even if their cell-level argmax differs only marginally. This rule enforces the principle that cell-level deviations from the population context are not trusted unless supported by strong individual evidence, thereby preventing scattered argmax errors from fragmenting otherwise coherent populations^35,36,37^.

#### Cells in Mixed clusters

For clusters labeled Mixed, representing populations with two competing lineages exhibiting comparable cluster-level support, MOSAIC adopts a conservative refinement strategy that preserves ambiguity at the cell level.

In these clusters, no cell-level identity is committed, regardless of individual probability differences, and all constituent cells are labeled Ambiguous. This design reflects the interpretation that, when population-level evidence does not support a single coherent lineage, cell-level deviations are not considered reliable indicators of definitive identity.

By refraining from forced disambiguation in Mixed clusters, MOSAIC explicitly preserves intermediate or transitional cellular states and avoids introducing spurious sharp boundaries within biologically heterogeneous populations.

#### Cells in Unknown clusters

For clusters labeled Unknown, marker evidence at the population level is deemed weak, inconsistent, or non-specific. In such cases, MOSAIC refrains from assigning cell-level identities, and all cells within these clusters are labeled Ambiguous.

This behavior reflects a deliberate design choice: when population-level evidence is insufficient to support a coherent lineage assignment, individual cell-level predictions are considered unreliable regardless of their probabilistic confidence^37,55^. By avoiding forced assignments in these contexts, MOSAIC prevents overinterpretation of poorly supported populations and preserves uncertainty as an explicit and meaningful outcome.

#### Final refined output

Taken together, the cell-level refinement module integrates cell-level probability profiles with cluster-level adaptive decisions, ensuring that final annotations are consistent with both local evidence and population-level structure. Rather than relying on cell-level argmax predictions in isolation, MOSAIC conditions individual cell identities on population context and explicitly retains ambiguity when available evidence is insufficient.

As a result, refined labels represent biologically coherent commitments, while uncertainty is preserved as a first-class outcome rather than treated as classification failure^35,36,37^.

### Method 6 | Benchmarking against external tools

To comprehensively evaluate performance across annotation paradigms, MOSAIC was benchmarked against five widely used cell-type annotation tools representing distinct methodological classes: SingleR^13^ (reference-based), CellTypist^15^ (large-scale supervised classification), scDeepSort^16^ (deep learning), and the marker-based approaches scCATCH^25^ and scType^11^.

All methods were executed using tissue-matched resources and author-recommended settings to ensure fair and method-compliant comparisons. SingleR was applied using the recommended human reference atlases under default settings^13^. CellTypist was run with tissue-appropriate pretrained models (e.g., immune, lung, pancreas) using the authors’ default inference parameters^15^. scDeepSort was evaluated using the published human pretrained weights, selecting tissue-matched models when available^16^.

Marker-based methods were applied using their respective tissue-specific marker databases and default scoring procedures. scType was executed following its original formulation, including direction-aware scoring based on positive and negative marker sets, without modification or additional parameter tuning^11^. The resulting scType score matrix was used for both cell-level and cluster-level evaluations.

As scCATCH produces cluster-level annotations only, its predicted cluster labels were propagated to constituent cells to obtain cell-level predictions^25^. For scType, cluster-level annotations were obtained using the standard scType workflow^11^, while cell-level predictions were obtained by assigning each cell to the highest-scoring cell type. For methods that natively produce cell-level outputs (SingleR, CellTypist, and scDeepSort), cluster-level predictions were derived by majority voting within Leiden clusters^24^ to enable direct comparison with the cluster-level outputs from MOSAIC.

All methods were evaluated at matched annotation granularity for each tissue using consistent label mappings and evaluation criteria, ensuring that observed performance differences reflect methodological capabilities rather than differences in resolution or task definition^10^.

### Method 7 | Evaluation metrics

Annotation performance was evaluated using a set of complementary metrics, with evaluation protocols tailored to the resolution of annotation (cell-level versus cluster-level) and to the availability of reliable ground-truth labels in different experimental settings^10^.

For real datasets, curated reference annotations were used as biological ground truth. Performance was evaluated separately at the cell level and the cluster level, reflecting the distinct roles of individual assignments and population-level coherence.

Cell-level performance was quantified using accuracy, defined as the fraction of cells whose predicted labels matched the curated reference annotations. For MOSAIC, cell-level predictions in the main text figures correspond to the initial soft cell-level label, obtained as the argmax of the per-cell probability vector derived from direction-aware marker scoring. This choice enables fair comparison with competing methods that produce a single deterministic label per cell.

In addition, MOSAIC provides an optional refined cell-level annotation, which assigns a definitive identity only when cell-level probabilistic evidence exceeds a predefined confidence threshold and is consistent with the adaptive cluster-level decision. Because this refined output represents selective commitment to high-confidence cells rather than a complete relabeling of all cells, its accuracy is reported separately in Supplementary Materials and is not used for primary cell-level comparisons in the main figures.

Cluster-level performance was evaluated using accuracy, computed by aggregating cell-level predictions within each Leiden cluster to obtain a single cluster label, which was then compared against the reference annotation. This metric quantifies the correctness of population-level assignments and is reported in Supplementary Materials. Cluster-level evaluation is particularly appropriate for real datasets, where transitional or ambiguous cells may exist and where population coherence is often the primary biological objective^24,54^.

For simulated datasets generated under controlled dropout perturbations, ground-truth labels are explicitly defined and preserved across all dropout levels. In this setting, performance was evaluated using a broader set of metrics to capture both classification accuracy and clustering consistency.

Specifically, Accuracy, Macro-F1, Adjusted Rand Index (ARI), and Normalized Mutual Information (NMI) were computed. Macro-F1 was calculated as the unweighted mean F1-score across cell types to mitigate class imbalance^56^. ARI measured agreement between predicted and ground-truth labels while correcting for chance^57^, and NMI quantified the mutual dependence between predicted and reference assignments normalized by their entropies^58^. Unless otherwise stated, these metrics were computed at the cluster level, consistent with the population-level nature of robustness analyses.

All metrics were computed using scikit-learn (v1.4.1). Accuracy, F1-score, ARI, and NMI were calculated using standard implementations without additional tuning^59^.

### Method 8 | Dropout simulation using scDesign2

To evaluate robustness under controlled technical sparsity, we started from baseline simulated count matrices generated using scDesign2 (v2.3), which were fitted to the corresponding real datasets in a separate step^27,28^. These scDesign2-simulated datasets served as the reference, unperturbed inputs.

First, using the filtered count matrix without additional dropout (dropout = 0), we applied a standard Scanpy preprocessing workflow, including cell and gene filtering, total-count normalization, log-transformation, highly variable gene selection, scaling, dimensionality reduction, and neighborhood graph construction^44^. Based on curated marker genes reported in the original studies, cell types were manually annotated on this reference dataset to obtain a unified set of ground-truth labels.

Next, to introduce controlled technical noise, additional dropout was applied directly to the filtered count matrix by randomly setting a specified fraction

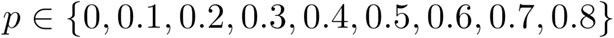

of non-zero entries to zero, while leaving existing zeros unchanged^27,28,60^. For each dropout level, the perturbed dataset was processed independently using the same normalization, transformation, feature selection, and dimensionality reduction pipeline.

All annotation methods were then applied separately to each dropout-perturbed dataset. Performance was evaluated using Accuracy, Macro-F1, ARI, and NMI by comparing predicted labels against the fixed ground-truth annotations derived from the dropout = 0 reference dataset. This design isolates the effect of increasing expression sparsity while keeping cell identities, marker definitions, and evaluation criteria constant.

Importantly, ground-truth annotations were defined once on the unperturbed reference dataset and were kept fixed across all dropout levels, preventing information leakage and ensuring that observed performance differences reflect robustness to increasing sparsity rather than changes in annotation criteria^28^.

### Method 9 | Visualization and implementation details

In selected figures, a permissive auxiliary probability threshold (*θ*_*vis*_ = 0.1) was applied to UMAP embeddings for visualization purposes only, to improve the visual interpretability of cells with weak or diffuse lineage support, including low-confidence and transitional states^44,45^. This auxiliary threshold was applied exclusively to PBMC visualizations in the main figures; all other tissues were visualized using default parameters without additional thresholding. Importantly, this threshold was not used for annotation, refinement, or benchmarking, and had no influence on any quantitative results or reported performance metrics. All algorithmic decisions, evaluations, and final labels were based exclusively on the probabilistic representations and thresholds defined in the corresponding method sections. An illustrative comparison of default and relaxed visualization thresholds is provided in Supplementary Fig. S9.

In line-plot visualizations comparing multiple annotation methods across discrete dropout levels, small fixed horizontal and vertical offsets were applied to individual curves for visualization purposes only, to reduce overplotting and improve visual separability^61^. These offsets were constant across dropout levels within each method and did not modify the underlying data values or relative method comparisons. All reported metrics, statistical summaries, and robustness scores were computed using the original, unshifted values.

## Competing interests

The authors declare no competing interests.

## Data availability

All datasets analyzed in this study are publicly available. PBMC data were obtained from 10x Genomics. Lung, liver, colon, pancreas, and retina datasets were obtained from publicly accessible repositories as detailed in the Methods. Processed data and intermediate results generated in this study will be made publicly available upon publication.

## Code availability

The MOSAIC framework was implemented in Python. The code and accompanying demonstration associated with this study have been deposited in a public GitHub repository: https://github.com/Mya-YANG/MOSAIC.

## Author contributions

**M.Y.** conceived the study, developed the MOSAIC framework, and performed all analyses. **S.J.** supervised the project. **M.Y.** wrote the manuscript with input from all authors.

---

**Extended Data Fig. 1.**
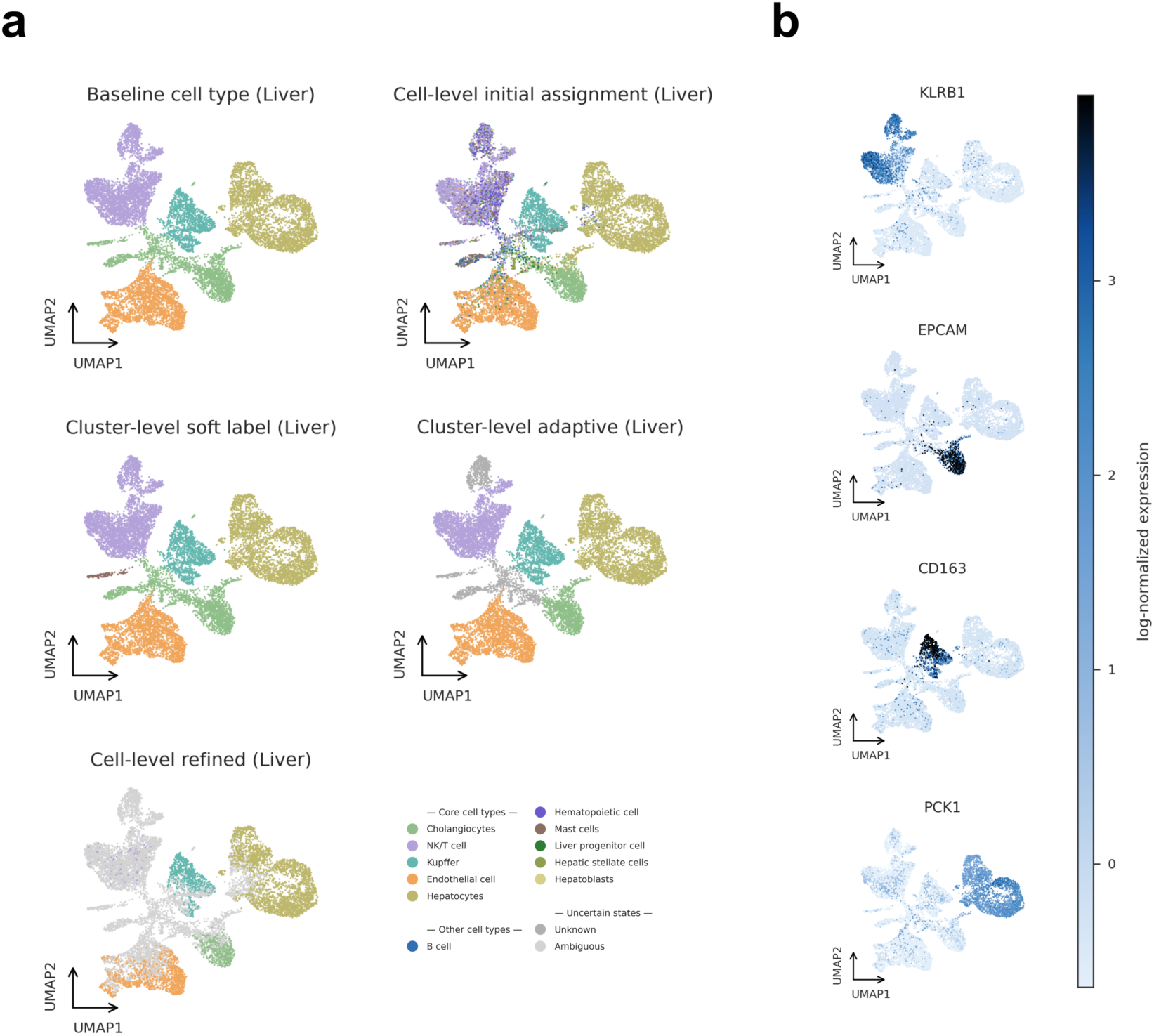
Multi-level probabilistic annotation behavior of MOSAIC in a non-immune tissue (liver). **(a)** UMAP visualizations showing successive stages of multi-level annotation by MOSAIC applied to liver tissue. Top left, curated baseline cell-type labels shown solely as a reference for visualization and evaluation. Top right, initial cell-level probabilistic assignments derived from direction-aware marker scoring, which recover major hepatic populations while exhibiting substantial local uncertainty, particularly at interfaces between transcriptionally related states. Middle left, cluster-level soft identities obtained by aggregating full cell-level probability distributions, revealing coherent population-level structure. Middle right, adaptive cluster-level decisions classifying clusters as Confident, Mixed, or Unknown based on internal agreement and overall support strength. Bottom, refined cell-level annotations obtained by reconciling cluster-level decisions with local probabilistic evidence, assigning labels only where consistent and retaining ambiguity otherwise. **(b)** UMAP overlays of representative hepatic marker genes confirm the inferred identities of major parenchymal and non-parenchymal populations and align with adaptively inferred cluster assignments. Together, this example demonstrates that probabilistic cell-level representations capture local uncertainty, whereas robustness and annotation stability emerge through population-level aggregation. These behaviors generalize beyond immune-centric systems, indicating that the multi-level inference principles underlying MOSAIC are applicable across diverse tissue contexts.

**Extended Data Fig. 2.**
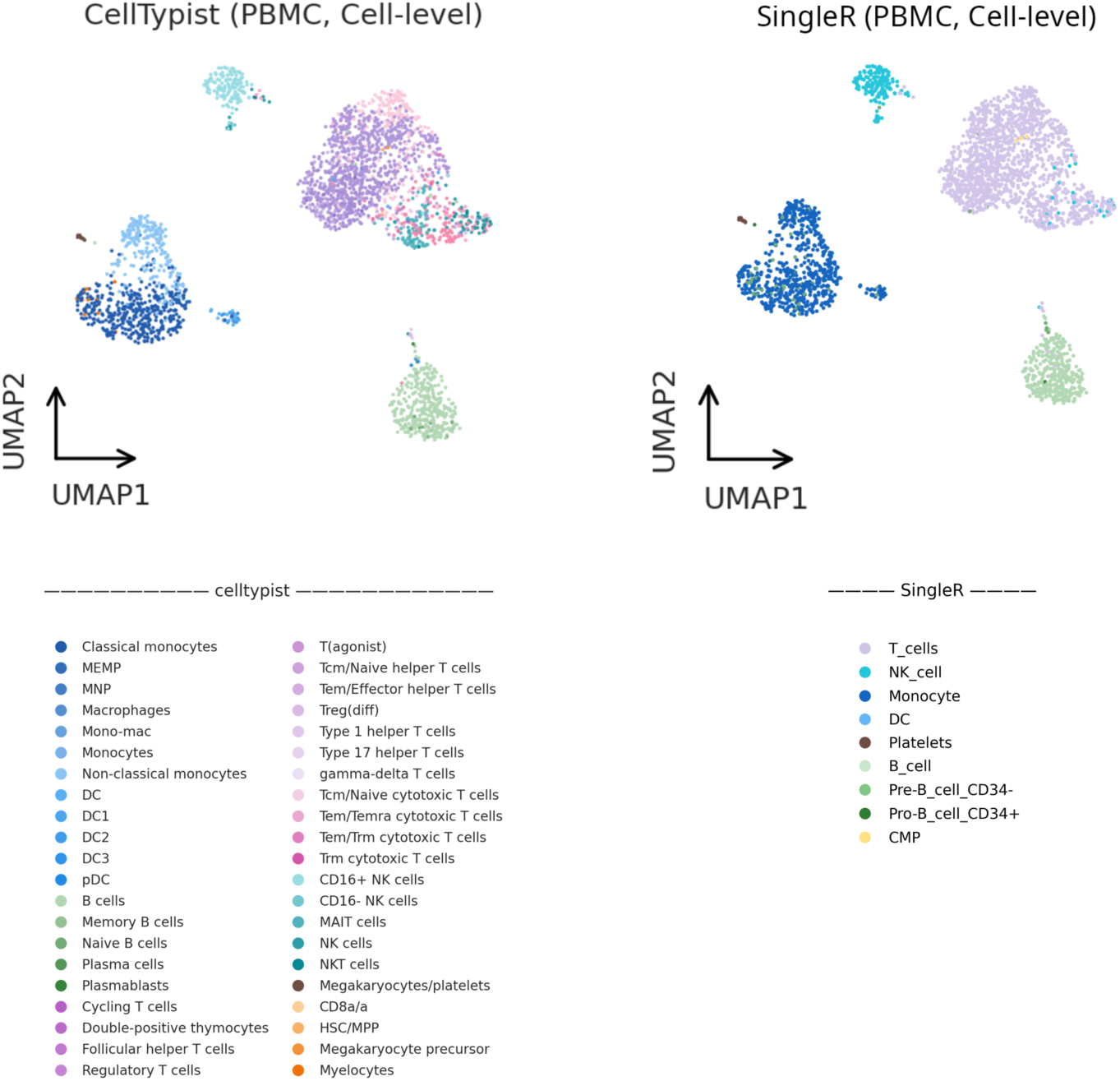
Deterministic annotation outputs of existing methods lack explicit uncertainty representation. UMAP visualizations of PBMC annotations produced by representative reference-based (SingleR) and large-scale classifier (CellTypist) methods at the cell level. Both approaches assign a single discrete label to each cell and do not provide an explicit probabilistic representation of uncertainty or confidence. As a result, annotation outputs appear uniformly resolved across the embedding, including regions corresponding to known lineage boundaries between closely related immune states. In some cases, composite or intermediate labels (for example, Tcm/Naive helper T cells in CellTypist) are used to capture biological relatedness. However, these identities are encoded as fixed categorical classes rather than emerging from graded probabilistic support. Consequently, ambiguity and transitional structure cannot be quantified, compared, or separated from confident assignments within a unified framework, limiting the ability to distinguish stable intermediate states from low-confidence predictions.

**Extended Data Fig. 3.**
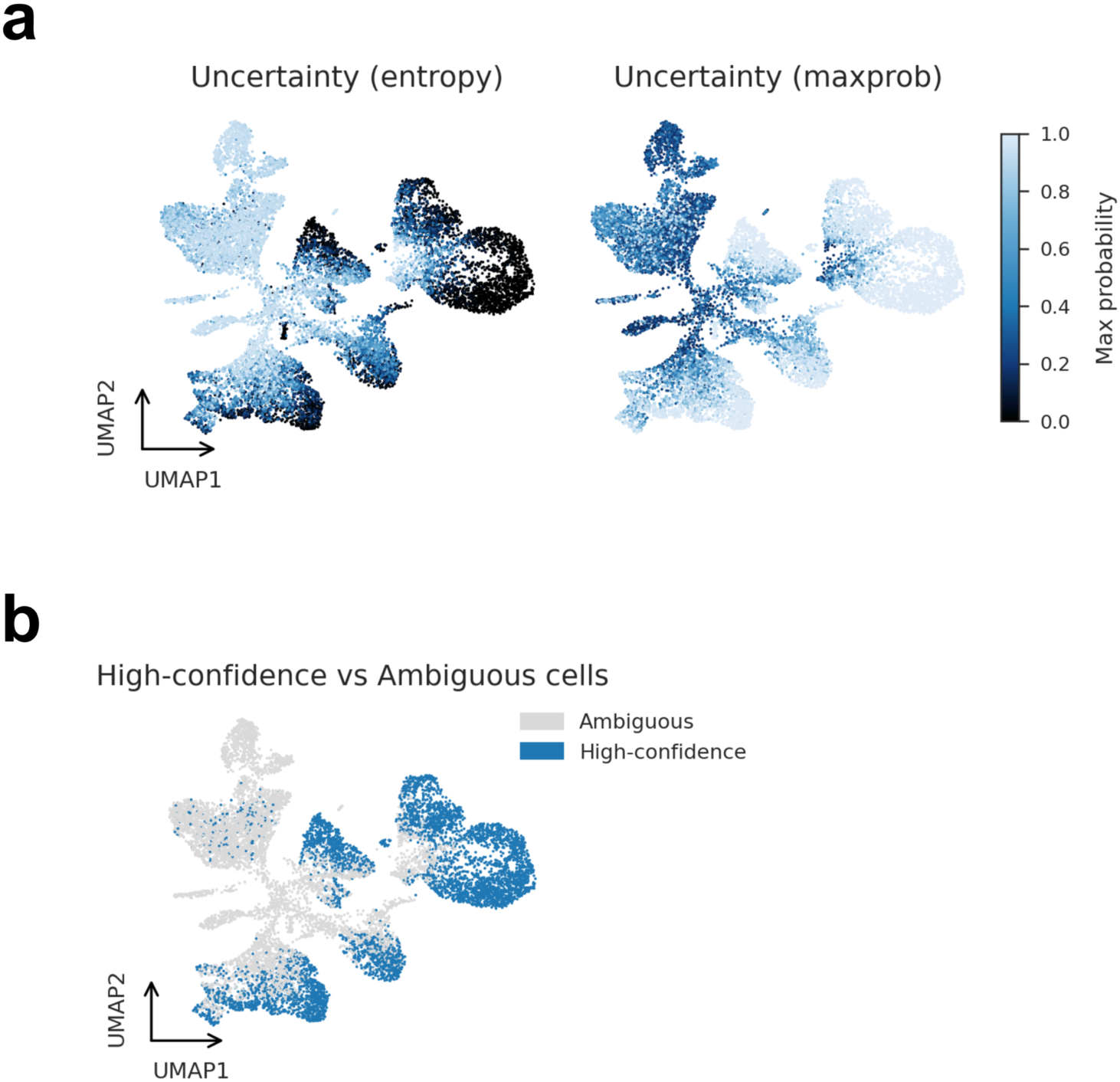
Structured cell-level uncertainty patterns generalize to liver tissue. **(a)** UMAP projections showing the spatial organization of cell-level uncertainty in liver tissue, quantified using normalized entropy and maximum class probability (maxprob) derived from full posterior probability vectors. Cells with elevated uncertainty preferentially localize to boundary regions between transcriptionally related hepatic lineages and to mixed cellular compartments, whereas well-separated populations exhibit uniformly low entropy and high maximum class probability. **(b)** Based on these uncertainty measures, cells can be categorized into high-confidence and ambiguous states without enforcing hard decisions. High-confidence cells form compact and biologically coherent populations, whereas ambiguous cells are enriched in transitional or weakly supported regions. Notably, these structured uncertainty patterns closely mirror those observed in PBMC, indicating that probabilistic ambiguity captured by MOSAIC reflects general properties of biological state transitions rather than tissue-specific artifacts.

## Notes

https://github.com/Mya-YANG/MOSAIC

### Competing Interest Statement

The authors have declared no competing interest.

