## Supplementary for "MOSAIC: A Structured Multi-level Framework for Probabilistic and Interpretable Cell-type Annotation"

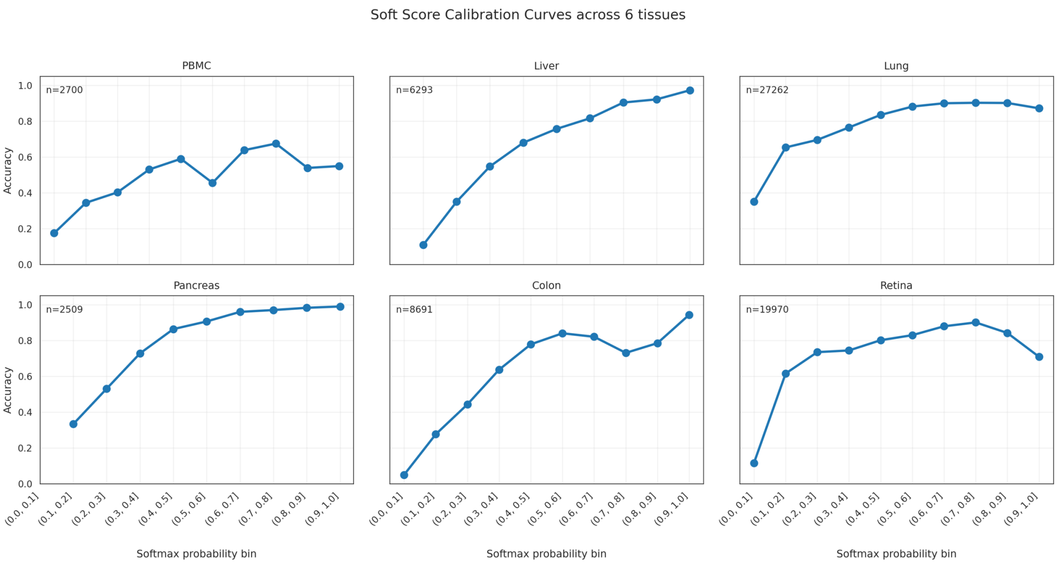


**Supplementary Fig. S1 | Calibration of probabilistic scores across tissues.**

Accuracy as a function of binned maximum softmax probability for PBMC, liver, lung, pancreas, colon, and retina. Higher probability bins generally correspond to increased annotation accuracy, indicating that probabilistic outputs produced by MOSAIC are well calibrated across diverse tissue contexts. Cell counts for each tissue are indicated. These curves demonstrate that probability values reflect meaningful confidence levels rather than arbitrary scores.

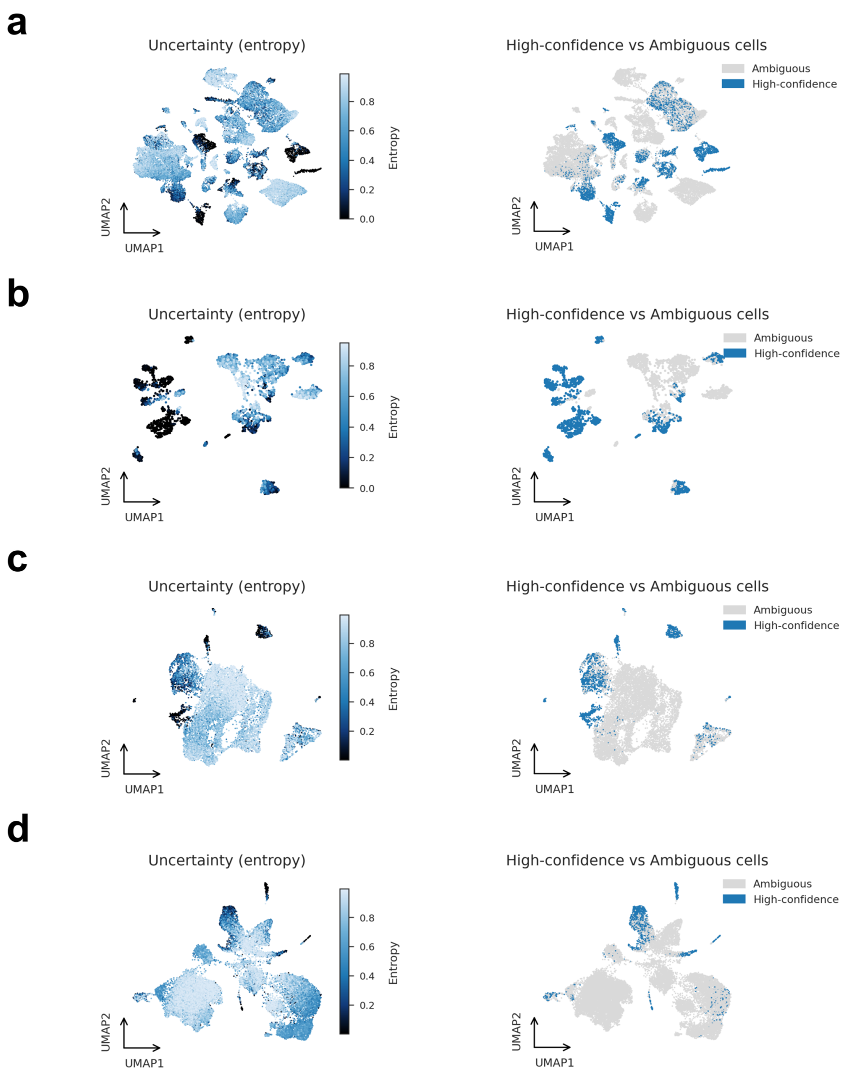


**Supplementary Fig. S2 | Uncertainty patterns across additional tissues.**

UMAP projections of normalized entropy and high-confidence versus ambiguous cells for additional tissues not shown in the main text: lung **(a)**, pancreas **(b)**, colon **(c)**, and retina **(d)**. Across all tissues examined, ambiguous cells tend to localize at interfaces between closely related lineages, whereas transcriptionally distinct populations are predominantly composed of high-confidence cells. These results indicate that structured uncertainty is consistently observed in MOSAIC outputs across diverse biological systems.

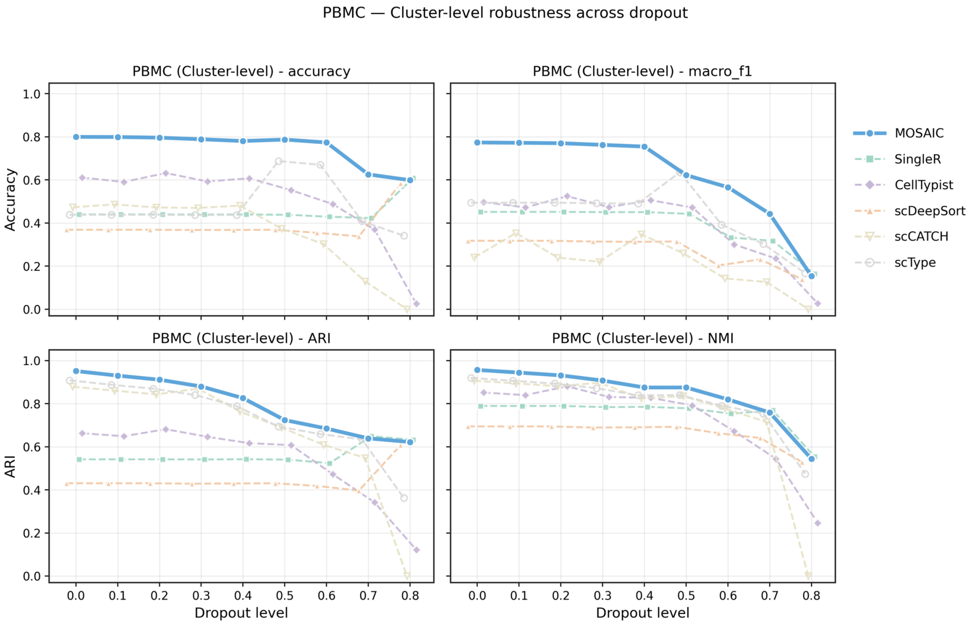


**Supplementary Fig. S3 | Cluster-level robustness under controlled dropout perturbations in PBMC**

Controlled dropout simulations were used to evaluate the robustness of cluster-level annotations in the PBMC dataset. Cluster-level accuracy, Macro-F1, ARI, and NMI were quantified across increasing dropout levels (0–0.8) and compared across representative annotation methods. Changes in performance under progressive marker loss are shown for each metric, illustrating the relative stability of population-level annotations produced by MOSAIC. AUC-based robustness summaries are provided in the corresponding Supplementary Tables.

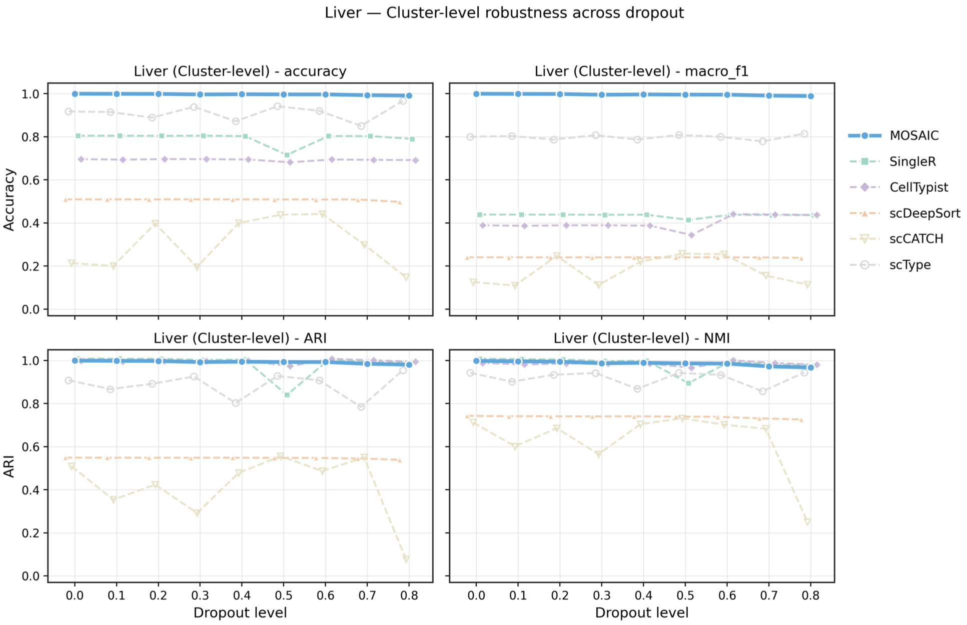


**Supplementary Fig. S4 | Cluster-level robustness under controlled dropout perturbations in liver**

Cluster-level robustness analysis for liver tissue under controlled dropout perturbations. Cluster-level accuracy, Macro-F1, ARI, and NMI were evaluated across increasing dropout levels (0–0.8) using scDesign2-based simulations. Results demonstrate that population-level annotations produced by MOSAIC remain stable under progressive expression loss.

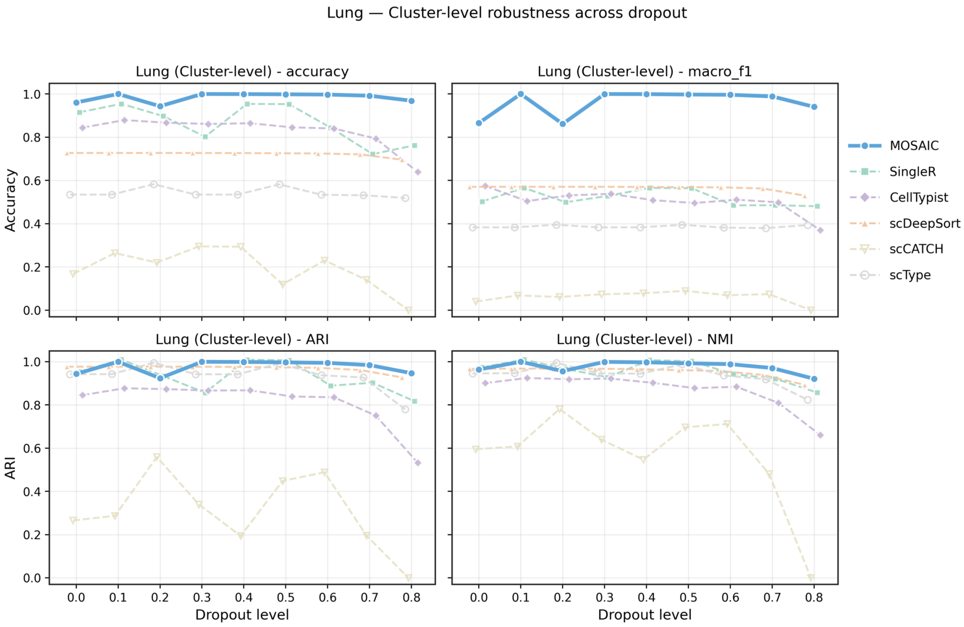


**Supplementary Fig. S5 | Cluster-level robustness under controlled dropout perturbations in lung**

Dropout robustness analysis of cluster-level annotations in lung tissue. Performance metrics including accuracy, Macro-F1, ARI, and NMI were assessed across increasing dropout levels (0–0.8) and compared across annotation methods. The resulting performance–dropout curves quantify the resilience of population-level structure to simulated marker loss.

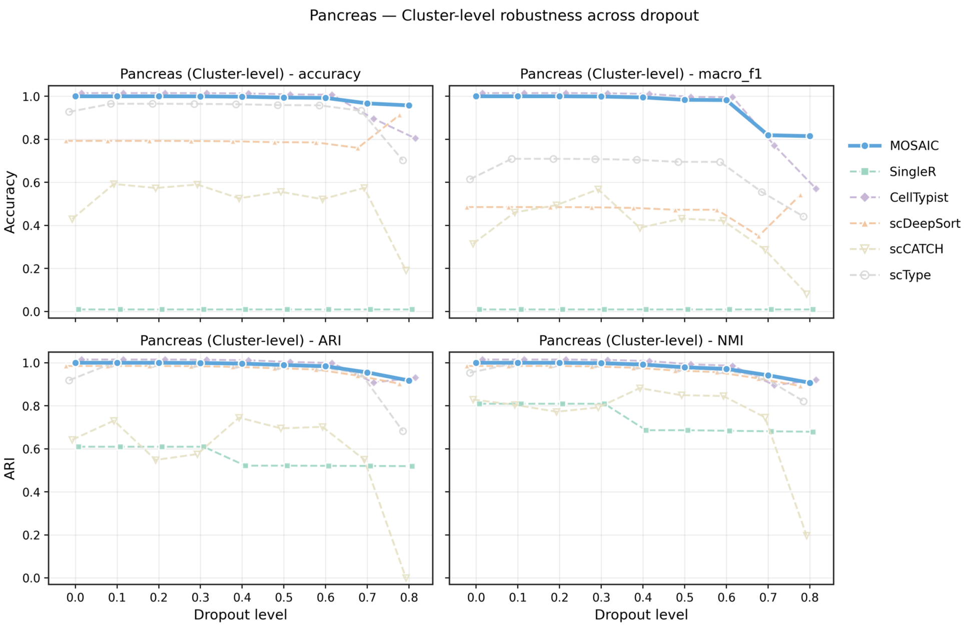


**Supplementary Fig. S6 | Cluster-level robustness under controlled dropout perturbations in pancreas**

Cluster-level robustness of annotations in pancreas under controlled dropout simulations. Accuracy, Macro-F1, ARI, and NMI were computed across increasing dropout levels. Pancreas, characterized by well-separated endocrine populations, serves as a reference context in which most methods achieve high baseline performance, enabling comparison of robustness trends under increasing perturbation.

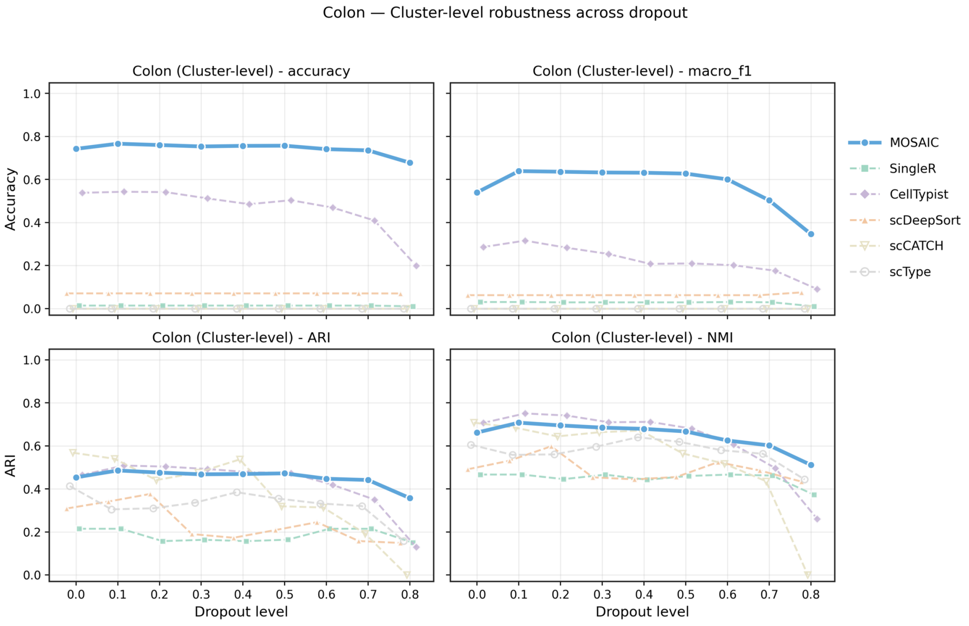


**Supplementary Fig. S7 | Cluster-level robustness under controlled dropout perturbations in colon**

Evaluation of cluster-level annotation robustness in colon tissue under simulated dropout perturbations. Cluster-level accuracy, Macro-F1, ARI, and NMI were quantified across dropout levels ranging from 0 to 0.8.

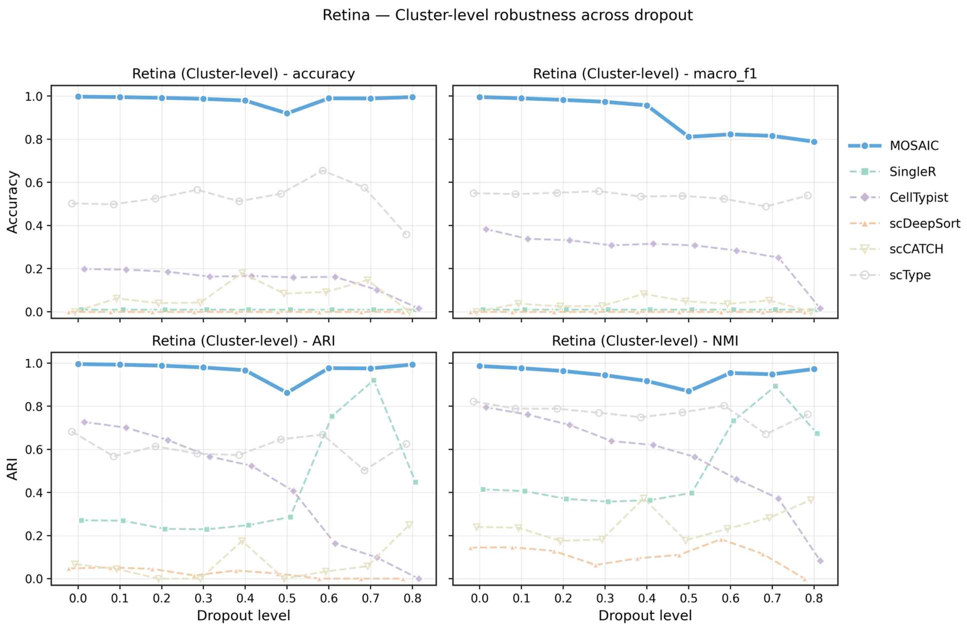


**Supplementary Fig. S8 | Cluster-level robustness under controlled dropout perturbations in retina**

Cluster-level robustness analysis for retina tissue under controlled dropout perturbations. Performance metrics including accuracy, Macro-F1, ARI, and NMI were assessed across increasing dropout levels.

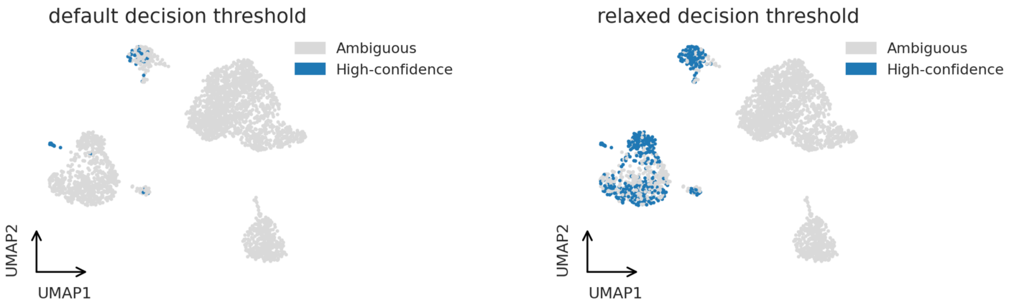


**Supplementary Fig. S9 | Effect of visualization-only cutoff on PBMC UMAP embeddings.**

Comparison of UMAP visualizations generated using the default cell-level commitment threshold ($\theta_{cell}$ = 0.8) and a relaxed auxiliary cutoff ($\theta_{vis}$ = 0.1) applied solely for visualization purposes. The relaxed cutoff increases the number of visually highlighted cells with weak or diffuse lineage support, facilitating inspection of low-confidence and transitional states.

Importantly, $\theta_{vis}$ is a display-only parameter and is distinct from the cell-level commitment threshold $\theta_{cell}$ used for annotation refinement. It does not alter probabilistic inference, cluster-level decisions, refined labels, or any quantitative evaluation. All reported results and benchmarks are based exclusively on $\theta_{cell}$ unless otherwise stated.

| tissue | MOSAIC | MOSAIC_refined | SingleR | CellTypist | scDeepSort | scCATCH | scType |
|---|---|---|---|---|---|---|---|
| **PBMC** | 0.5111 | 1.0 | 0.1922 | 0.4189 | 0.1411 | 0.2811 | 0.3807 |
| **Liver** | 0.7703 | 1.0 | 0.6971 | 0.5665 | 0.468 | 0.3347 | 0.5443 |
| **Colon** | 0.6809 | 1.0 | 0.0 | 0.6798 | 0.1031 | 0.0 | 0.3981 |
| **Pancreas** | 0.9095 | 1.0 | 0.0 | 0.9378 | 0.7991 | 0.6584 | 0.6903 |
| **Lung** | 0.8251 | 1.0 | 0.7893 | 0.7436 | 0.6467 | 0.388 | 0.4017 |
| **Retina** | 0.7643 | 0.9419 | 0.0 | 0.1544 | 0.0 | 0.0746 | 0.2459 |

**Supplementary Table S1 | Cell-level annotation accuracy on real datasets**

Cell-level annotation accuracy across multiple real scRNA-seq datasets, including PBMC and non-immune tissues. Accuracy is reported for MOSAIC (initial and refined) and competing annotation methods. Cell-level performance is summarized using accuracy, as clustering-oriented metrics are more appropriately evaluated at the population (cluster) level.

| tissue | MOSAIC | SingleR | CellTypist | scDeepSort | scCATCH | scType |
|---|---|---|---|---|---|---|
| **PBMC** | 1.0 | 0.1937 | 0.407 | 0.1459 | 0.2811 | 0.6481 |
| **Liver** | 0.9891 | 0.7351 | 0.5768 | 0.4782 | 0.3347 | 0.8458 |
| **Colon** | 1.0 | 0.0 | 0.7257 | 0.1052 | 0.0 | 0.5739 |
| **Pancreas** | 1.0 | 0.0 | 0.9817 | 0.8183 | 0.6584 | 0.9028 |
| **Lung** | 0.9859 | 0.9411 | 0.7971 | 0.7111 | 0.388 | 0.5958 |
| **Retina** | 0.994 | 0.0 | 0.1635 | 0.0 | 0.0746 | 0.6141 |

**Supplementary Table S2 | Cluster-level annotation accuracy on real datasets**

Cluster-level annotation accuracy across real scRNA-seq datasets. For each tissue, cell-level predictions are aggregated to derive cluster-level labels, which are then compared against curated reference annotations. Accuracy quantifies the correctness of population-level assignments across methods.

| tissue | MOSAIC | SingleR | CellTypist | scDeepSort | scCATCH | scType |
|---|---|---|---|---|---|---|
| **PBMC** | 1.0 | 0.1818 | 0.3262 | 0.1818 | 0.2308 | 0.7187 |
| **Liver** | 0.8275 | 0.3567 | 0.1744 | 0.2148 | 0.125 | 0.7332 |
| **Colon** | 1.0 | 0.0 | 0.3484 | 0.1111 | 0.0 | 0.1789 |
| **Pancreas** | 1.0 | 0.0 | 0.7378 | 0.5556 | 0.4241 | 0.6486 |
| **Lung** | 0.9867 | 0.4967 | 0.3536 | 0.4461 | 0.0874 | 0.3636 |
| **Retina** | 0.6364 | 0.0 | 0.2208 | 0.0 | 0.0368 | 0.4862 |

**Supplementary Table S3 | Cluster-level Macro-F1 scores on real datasets**

Cluster-level Macro-F1 scores across real scRNA-seq datasets. Macro-F1 accounts for class imbalance by equally weighting cell-type categories and highlights balanced population-level annotation performance across both major and minor cell types. Scores are reported after aggregating cell-level predictions to the cluster level.

| tissue | method | ARI | NMI |
|---|---|---|---|
| **PBMC** | MOSAIC | 1.0 | 1.0 |
| **PBMC** | SingleR | 0.4588801427674470 | 0.7336173396977620 |
| **PBMC** | CellTypist | 0.5752358295097380 | 0.7521370683556080 |
| **PBMC** | scDeepSort | 0.3780359818202770 | 0.6694296754814120 |
| **PBMC** | scCATCH | 0.8401565755812630 | 0.9197092978040870 |
| **PBMC** | scType | 0.9420511630603100 | 0.9678840035683300 |
| **Liver** | MOSAIC | 0.9903757067080120 | 0.9873402344450690 |
| **Liver** | SingleR | 0.8346859391430370 | 0.8897503065966800 |
| **Liver** | CellTypist | 0.8122764514971850 | 0.8190126326807370 |
| **Liver** | scDeepSort | 0.481577083670912 | 0.7203637972128870 |
| **Liver** | scCATCH | 0.6081848324370040 | 0.7806780552032650 |
| **Liver** | scType | 0.7895588381963410 | 0.8505600198342140 |
| **Colon** | MOSAIC | 1.0 | 1.0 |
| **Colon** | SingleR | 0.3998796291041880 | 0.537801968745287 |
| **Colon** | CellTypist | 0.7963762746945540 | 0.7980347097192130 |
| **Colon** | scDeepSort | 0.5201062598443380 | 0.7185905602028550 |
| **Colon** | scCATCH | 0.2278181465375770 | 0.5157306011078380 |
| **Colon** | scType | 0.5317039073529080 | 0.7567414140594680 |
| **Pancreas** | MOSAIC | 1.0 | 1.0 |
| **Pancreas** | SingleR | 0.3294639319224550 | 0.5513031555968150 |
| **Pancreas** | CellTypist | 0.9736046071541970 | 0.9653348411391960 |
| **Pancreas** | scDeepSort | 1.0 | 1.0 |
| **Pancreas** | scCATCH | 0.6772048648609390 | 0.8114605248892930 |
| **Pancreas** | scType | 0.867748926898854 | 0.9081096008312060 |
| **Lung** | MOSAIC | 0.9753297887677700 | 0.9701841355163940 |
| **Lung** | SingleR | 0.969778552240402 | 0.9763857472464560 |
| **Lung** | CellTypist | 0.7160352132866230 | 0.7652502119640770 |

| | | | |
|---|---|---|---|
| **Lung** | scDeepSort | 0.9537913419674610 | 0.9308232756829790 |
| **Lung** | scCATCH | 0.7198846776269590 | 0.8198876356501900 |
| **Lung** | scType | 0.9014220886491210 | 0.925979167891304 |
| **Retina** | MOSAIC | 1.0 | 1.0 |
| **Retina** | SingleR | 0.4147095926512640 | 0.5340675790092840 |
| **Retina** | CellTypist | 0.6991461105721700 | 0.7601580046303150 |
| **Retina** | scDeepSort | 0.119365015102361 | 0.2544962730057920 |
| **Retina** | scCATCH | 0.1451584803285510 | 0.3335031631273370 |
| **Retina** | scType | 0.7072811908374270 | 0.8417068282576190 |

**Supplementary Table S4 | Cluster-level ARI and NMI on real datasets**

Adjusted Rand Index (ARI) and Normalized Mutual Information (NMI) for cluster-level annotations across real scRNA-seq datasets. These metrics evaluate the agreement between predicted and reference population-level label partitions, assessing the preservation of population structure across methods.

| | MOSAIC | SingleR | CellTypist | scDeepSort | scCATCH | scType |
|---|---|---|---|---|---|---|
| **0.0** | 0.7997038134024440 | 0.4298407997038130 | 0.5957052943354310 | 0.3835616438356160 | 0.4827841540170310 | 0.4431691965938540 |
| **0.1** | 0.7989633469085520 | 0.4298407997038130 | 0.5753424657534250 | 0.3835616438356160 | 0.4961125509070710 | 0.4431691965938540 |
| **0.2** | 0.7960014809329880 | 0.4298407997038130 | 0.6168085894113290 | 0.3835616438356160 | 0.4820436875231400 | 0.4431691965938540 |
| **0.3** | 0.7885968159940760 | 0.4294705664568680 | 0.5775638652350980 | 0.3824509440947800 | 0.4798222880414660 | 0.4420584968530170 |
| **0.4** | 0.7802889959244170 | 0.429418303075213 | 0.5924416450537240 | 0.3827343460540940 | 0.4905520563171550 | 0.4423860689144130 |
| **0.5** | 0.7871709306637000 | 0.4278828327771600 | 0.5372636262513910 | 0.3833889506859470 | 0.3833889506859470 | 0.6922506488691140 |
| **0.6** | 0.7735217552993680 | 0.4187430271476390 | 0.4726664187430270 | 0.3692822610635920 | 0.3123837857939750 | 0.6753439940498330 |
| **0.7** | 0.6254927726675430 | 0.4130530004380200 | 0.3543583004818220 | 0.3526062198861150 | 0.1388523872098120 | 0.4130530004380200 |
| **0.8** | 0.5986159169550170 | 0.5986159169550170 | 0.0103806228373702 | 0.5986159169550170 | 0.0 | 0.3460207612456750 |
| **AUC** | 0.7561494954461720 | 0.4365597109539930 | 0.503685983689527 | 0.3910843487313850 | 0.3780684729358850 | 0.4932531976539840 |

**Supplementary Table S5 | Cluster-level accuracy across dropout levels in PBMC**

Cluster-level annotation accuracy under increasing scDesign2-simulated dropout levels in the PBMC dataset. Accuracy is reported for dropout fractions ranging from 0 to 0.8, together with the area under the accuracy–dropout curve (AUC) as an aggregate robustness measure. This table summarizes the performance of population-level annotations produced by MOSAIC and competing methods under progressive marker loss.

|  | MOSAIC | SingleR | CellTypist | scDeepSort | scCATCH | scType |
|---|---|---|---|---|---|---|
| **0.0** | 0.7736663301443840 | 0.4414504324683970 | 0.4817934987925980 | 0.3322332233223320 | 0.2497267759562840 | 0.4990771502399410 |
| **0.1** | 0.7720518248605620 | 0.4414504324683970 | 0.4570769980295680 | 0.3322332233223320 | 0.3634870876250190 | 0.4991680532445920 |
| **0.2** | 0.7701942564564490 | 0.4414504324683970 | 0.5095770487393600 | 0.3322332233223320 | 0.2495881383855020 | 0.499350769801521 |
| **0.3** | 0.7624789777386950 | 0.4402501332643000 | 0.4583429802810330 | 0.3288919000301110 | 0.2296986082791470 | 0.4960995589991140 |
| **0.4** | 0.7546233509689940 | 0.4403356052666130 | 0.4906271763917080 | 0.3274528607962140 | 0.357301357473484 | 0.4954698265707670 |
| **0.5** | 0.6222803298213570 | 0.4322323558407700 | 0.4564759298695700 | 0.3292264477619170 | 0.2699205629366800 | 0.6405223775778260 |
| **0.6** | 0.5658782320740190 | 0.3226185805572450 | 0.2852153805528450 | 0.2181819884881440 | 0.152631889615531 | 0.3971823211720280 |
| **0.7** | 0.4429018977280480 | 0.3072411472670090 | 0.2197250153884530 | 0.2460709345010510 | 0.136129233136138 | 0.3072411472670090 |
| **0.8** | 0.1533687943262410 | 0.1533687943262410 | 0.0115384615384615 | 0.1533687943262410 | 0.0 | 0.1720366421858960 |
| **AUC** | 0.6442408039854300 | 0.390373537566256 | 0.3904633136772580 | 0.2946364483807990 | 0.2354525331787050 | 0.4588238688557220 |

**Supplementary Table S6 | Cluster-level Macro-F1 across dropout levels in PBMC**

Cluster-level Macro-F1 scores under increasing scDesign2-simulated dropout levels in the PBMC dataset. Macro-F1 is reported for dropout fractions ranging from 0 to 0.8, together with the area under the Macro-F1–dropout curve (AUC) as an aggregate robustness measure. This table summarizes changes in balanced population-level annotation performance for MOSAIC and competing methods under progressive marker loss.

| | MOSAIC | SingleR | CellTypist | scDeepSort | scCATCH | scType |
|---|---|---|---|---|---|---|
| **0.0** | 0.9514645095314570 | 0.5316070196540800 | 0.6478240427848610 | 0.4455138075280100 | 0.8887860697989870 | 0.913391899887816 |
| **0.1** | 0.9301561423310220 | 0.5316070196540800 | 0.633893232722031 | 0.4455138075280100 | 0.8705406945674270 | 0.8927980185794680 |
| **0.2** | 0.9113023712184150 | 0.5316070196540800 | 0.66657426162508 | 0.4455138075280100 | 0.853045256724395 | 0.8746730375640460 |
| **0.3** | 0.879850708908395 | 0.5311009174499740 | 0.6311987008737970 | 0.4435795179758020 | 0.879850708908395 | 0.8452997924190680 |
| **0.4** | 0.8257850501398960 | 0.5320422940912990 | 0.6017183161415770 | 0.4446281531670960 | 0.7737022077252980 | 0.793356017130878 |
| **0.5** | 0.7239509716208100 | 0.530363592047894 | 0.5925910772836530 | 0.4453399150042860 | 0.6989240737172820 | 0.6989240737172820 |
| **0.6** | 0.6846439539249730 | 0.5137594286577490 | 0.4575188177903420 | 0.4331718129759050 | 0.619319127001982 | 0.6640345476615790 |
| **0.7** | 0.6390806978593070 | 0.6390806978593070 | 0.3266714391515660 | 0.4115525731492360 | 0.5584267592208270 | 0.6390806978593070 |
| **0.8** | 0.6222173222224620 | 0.6222173222224620 | 0.1069334147058130 | 0.6222173222224620 | 0.0 | 0.3674301436140450 |
| **AUC** | 0.7977013514849720 | 0.5483091425440820 | 0.5359430717916730 | 0.4503956440254480 | 0.7122752328456370 | 0.7560721508353200 |

**Supplementary Table S7 | Cluster-level ARI across dropout levels in PBMC**

Adjusted Rand Index (ARI) for cluster-level annotations in the PBMC dataset under increasing scDesign2-simulated dropout levels. ARI is reported across dropout fractions from 0 to 0.8, together with the area under the ARI–dropout curve (AUC) as an aggregate robustness measure.

| | MOSAIC | SingleR | CellTypist | scDeepSort | scCATCH | scType |
|---|---|---|---|---|---|---|
| **0.0** | 0.956564130618291 | 0.779234971764241 | 0.8365909431944140 | 0.709481236568096 | 0.9165389593185050 | 0.9241352439908320 |
| **0.1** | 0.9442266617045800 | 0.779234971764241 | 0.8247512686351620 | 0.709481236568096 | 0.9042547441992460 | 0.911437305490652 |
| **0.2** | 0.9312376543002900 | 0.779234971764241 | 0.8655914818874890 | 0.709481236568096 | 0.8909179000534910 | 0.8981270979301280 |
| **0.3** | 0.9073492292398230 | 0.7739581356777740 | 0.8162099075594440 | 0.7040225153051490 | 0.9073492292398230 | 0.8769397962906660 |
| **0.4** | 0.8751100132296040 | 0.7753748770679130 | 0.8121444838515350 | 0.705385059070863 | 0.8345003144597330 | 0.8438452312386500 |
| **0.5** | 0.8752045457712660 | 0.7682521955162000 | 0.7768327261861350 | 0.7074828450522740 | 0.844976540210645 | 0.844976540210645 |
| **0.6** | 0.8200692463304030 | 0.7432723959258850 | 0.6573529888479140 | 0.6779553942707580 | 0.7878365115859740 | 0.7952357258029850 |
| **0.7** | 0.7590841136199510 | 0.7590841136199510 | 0.5293797125815290 | 0.6554553410106210 | 0.724898392358974 | 0.7590841136199510 |
| **0.8** | 0.5438513711327130 | 0.5438513711327130 | 0.2315921438328700 | 0.5438513711327130 | 0.0 | 0.4793632455063730 |
| **AUC** | 0.8578111518839270 | 0.7549943540980850 | 0.7270442641328560 | 0.6869912414620330 | 0.7941253889708920 | 0.8289243819165350 |

**Supplementary Table S8 | Cluster-level NMI across dropout levels in PBMC**

Normalized Mutual Information (NMI) for cluster-level annotations in the PBMC dataset under increasing scDesign2-simulated dropout levels. NMI is reported across dropout fractions from 0 to 0.8, together with the area under the NMI–dropout curve (AUC) as an aggregate robustness measure.

| tissue | method | Acc-AUC | Macro-F1 AUC | ARI-AUC | NMI-AUC |
|---|---|---|---|---|---|
| **PBMC** | MOSAIC | 0.756149 | 0.644241 | 0.797701 | 0.857811 |
| **PBMC** | SingleR | 0.43656 | 0.390374 | 0.548309 | 0.754994 |
| **PBMC** | CellTypist | 0.503686 | 0.390463 | 0.535943 | 0.727044 |
| **PBMC** | scDeepSort | 0.391084 | 0.294636 | 0.450396 | 0.686991 |
| **PBMC** | scCATCH | 0.378068 | 0.235453 | 0.712275 | 0.794125 |
| **PBMC** | scType | 0.493253 | 0.458824 | 0.756072 | 0.828924 |
| **Lung** | MOSAIC | 0.986841 | 0.968244 | 0.980351 | 0.980779 |
| **Lung** | SingleR | 0.86071 | 0.512946 | 0.926604 | 0.951611 |
| **Lung** | CellTypist | 0.821622 | 0.492314 | 0.809714 | 0.862372 |
| **Lung** | scDeepSort | 0.738745 | 0.5814 | 0.985764 | 0.971877 |
| **Lung** | scCATCH | 0.214904 | 0.075693 | 0.340015 | 0.604592 |
| **Lung** | scType | 0.54991 | 0.3909 | 0.947209 | 0.949762 |
| **Liver** | MOSAIC | 0.996704 | 0.995675 | 0.992896 | 0.986815 |
| **Liver** | SingleR | 0.782118 | 0.4247 | 0.972562 | 0.974112 |
| **Liver** | CellTypist | 0.677813 | 0.383449 | 0.980654 | 0.96965 |
| **Liver** | scDeepSort | 0.523352 | 0.255029 | 0.562386 | 0.75346 |
| **Liver** | scCATCH | 0.328868 | 0.194255 | 0.439091 | 0.654199 |
| **Liver** | scType | 0.913647 | 0.802315 | 0.884819 | 0.920206 |
| **Colon** | MOSAIC | 0.747367 | 0.588977 | 0.458077 | 0.655981 |
| **Colon** | SingleR | 0.003335 | 0.017903 | 0.173434 | 0.443661 |
| **Colon** | CellTypist | 0.463824 | 0.214084 | 0.42519 | 0.632467 |
| **Colon** | scDeepSort | 0.085184 | 0.077722 | 0.255026 | 0.509375 |
| **Colon** | scCATCH | 0.0 | 0.0 | 0.397224 | 0.576254 |
| **Colon** | scType | 0.0 | 0.0 | 0.33351 | 0.585137 |
| **Pancreas** | MOSAIC | 0.991268 | 0.960737 | 0.985357 | 0.979529 |
| **Pancreas** | SingleR | 0.0 | 0.0 | 0.54986 | 0.728871 |

| | | | | | |
|---|---|---|---|---|---|
| **Pancreas** | CellTypist | 0.969976 | 0.93659 | 0.97746 | 0.971751 |
| **Pancreas** | scDeepSort | 0.809238 | 0.482991 | 0.985357 | 0.979529 |
| **Pancreas** | scCATCH | 0.540448 | 0.415964 | 0.617393 | 0.785666 |
| **Pancreas** | scType | 0.945423 | 0.668075 | 0.966234 | 0.971905 |
| **Retina** | MOSAIC | 0.981074 | 0.905462 | 0.967058 | 0.943975 |
| **Retina** | SingleR | 0.0 | 0.0 | 0.402206 | 0.498239 |
| **Retina** | CellTypist | 0.139546 | 0.276529 | 0.411171 | 0.556135 |
| **Retina** | scDeepSort | 0.0 | 0.0 | 0.03256 | 0.128105 |
| **Retina** | scCATCH | 0.089248 | 0.047107 | 0.060067 | 0.255032 |
| **Retina** | scType | 0.543468 | 0.540375 | 0.605861 | 0.771656 |

**Supplementary Table S9 | Summary of dropout robustness across tissues and methods (AUC)**

Area under the dropout–performance curves (AUC) summarizing robustness across tissues and annotation methods. For each tissue, AUC values are computed from cluster-level performance curves across increasing scDesign2-simulated dropout levels (0–0.8), including Accuracy, Macro-F1, ARI, and NMI.

| diff_leiden_resolution | Non-ambiguous (n) | Non-ambiguous (%) | Accuracy | Macro-F1 | ARI | NMI | n_total_cells |
|---|---|---|---|---|---|---|---|
| **leiden resolution=0.1** | 29 | 1.1 | 1.000 | 1.000 | 1.000 | 1.000 | 2700 |
| **leiden resolution=0.3** | 45 | 1.7 | 1.000 | 1.000 | 1.000 | 1.000 | 2700 |
| **leiden resolution=0.5** | 32 | 1.2 | 1.000 | 1.000 | 1.000 | 1.000 | 2700 |
| **(default) leiden resolution=0.6** | 45 | 1.7 | 1.000 | 1.000 | 1.000 | 1.000 | 2700 |
| **leiden resolution=0.8** | 47 | 1.7 | 0.957 | 0.786 | 0.950 | 0.941 | 2700 |

**Supplementary Table S10 | Sensitivity of refined annotation to Leiden clustering resolution**

Sensitivity analysis of MOSAIC refined annotations under varying Leiden clustering resolutions. Leiden resolution was varied from 0.1 to 0.8 while keeping all MOSAIC scoring, aggregation, and refinement procedures fixed. Metrics were computed on non-ambiguous cells retained after refinement, including cell-level Accuracy and Macro-F1, as well as cluster-level agreement with ground truth quantified by Adjusted Rand Index (ARI) and Normalized Mutual Information (NMI).

Across a broad and practically relevant range of resolutions (0.3–0.6), MOSAIC exhibits stable performance, maintaining consistently high accuracy and near-perfect cluster-level concordance. At very low resolution, refinement is conservative and restricted to a small number of high-confidence cells, whereas overly fine resolutions lead to mild performance degradation due to cluster over-fragmentation. Overall, these results demonstrate that MOSAIC's multi-level inference is robust to clustering granularity and does not depend on fine-tuned Leiden resolution parameters.

| Refined prob. threshold | Non-ambiguous (n) | Non-ambiguous (%) | Accuracy | Macro-F1 | ARI | NMI |
|---|---|---|---|---|---|---|
| **default (θ = 0.8)** | 45 | 1.7 | 1.0000 | 1.0000 | 1.000 | 1.000 |
| **0.7** | 75 | 2.8 | 1.0000 | 1.0000 | 1.000 | 1.000 |
| **0.5** | 139 | 5.1 | 1.0000 | 1.0000 | 1.000 | 1.000 |
| **0.3** | 304 | 11.3 | 1.0000 | 1.0000 | 1.000 | 1.000 |
| **0.1** | 480 | 17.8 | 1.0000 | 1.0000 | 1.000 | 1.000 |
| **0** | 483 | 17.9 | 1.0000 | 1.0000 | 1.000 | 1.000 |

**Supplementary Table S11 | Sensitivity of refined cell-level commitment thresholds in PBMC.**

Summary of annotation outcomes under varying cell-level commitment thresholds ($\theta_{cell}$). Lower thresholds increase the fraction of cells labeled Confident, while overall accuracy, Macro-F1, ARI, and NMI remain stable. These results indicate that $\theta_{cell}$ primarily controls label commitment stringency rather than affecting overall annotation performance.
